# Intrinsically disordered region amplifies membrane remodeling to augment selective ER-phagy

**DOI:** 10.1101/2024.03.28.587138

**Authors:** Sergio Alejandro Poveda-Cuevas, Kateryna Lohachova, Ivan Dikic, Gerhard Hummer, Ramachandra M. Bhaskara

## Abstract

Intrinsically disordered regions (IDRs) play a pivotal role in organellar remodeling. They transduce signals across membranes, scaffold signaling complexes, and mediate vesicular traffic. Their functions are regulated by constraining conformational ensembles through specific intra- and inter-molecular interactions, physical tethering, and post-translational modifications. The ER-phagy receptor FAM134B/RETREG1, known for its Reticulon homology domain (RHD), includes a substantial C-terminal intrinsically disorder region (IDR) housing the LC3 interacting (LIR) motif. Beyond engaging the autophagic machinery, the function of the FAM134B-IDR is unclear. Here, we investigate the characteristics of the FAM134B-IDR by extensive modeling and molecular dynamics (MD) simulations. We present detailed structural models for the IDR, mapping its conformational landscape in solution and membrane-anchored configurations. Our analysis reveals that depending on the membrane anchor, the IDRs collapse onto the membrane and induce positive membrane curvature to varying degrees. The charge patterns underlying this Janus-like behavior are conserved across other ER-phagy receptors. We found that IDRs alone are sufficient to sense curvature. When combined with RHDs, they intensify membrane remodeling and drive efficient protein clustering, leading to faster budding, thereby amplifying RHD remodeling functions. Our simulations provide a new perspective on IDRs of FAM134B, their Janus-like membrane interactions, and the resulting modulatory functions during large-scale ER remodeling.

## I. INTRODUCTION

Exploring the human proteome reveals a complex and dynamic landscape woven into an intricate protein network that governs a diverse array of cellular processes. A significant subset contains intrinsically disordered regions (IDRs), segments that lack a stable 3D structure under physiological conditions (Van Der Lee et al. 2014). The diversity of IDRs within the human proteome plays a pivotal role in regulating a vast array of cellular functions (Cornish et al. 2020). From facilitating protein-protein interactions (PPIs) and signal transduction to orchestrating transcriptional and post-translational modifications (PTMs), IDRs serve as versatile modules contributing to the complexity and adaptability of the cellular machinery (Van Der Lee et al. 2014). Further, multivalent interactions of the IDRs promote liquid-liquid phase separation (LLPS) to form condensates, which concentrate specific proteins and nucleic acids for localized cellular action (Tesei et al. 2021).

Membrane-associated and membrane-anchored IDRs have emerged as dynamic mediators of processes occurring at the surface of membrane-bound organelles (Cornish et al. 2020). IDRs adopt diverse biochemical and biophysical principles to actively bind and modulate membrane functions, *e.g*., by folding of disordered regions upon membrane binding (disorder-to-order transitions) and by forming amphipathic helices that induce or sense membrane curvature (Cornish et al. 2020). 2D confinement of IDRs to membranes increases their effective concentration and limits the search space for IDR-mediated PPIs, thereby enhancing protein association and crowding (Stachowiak et al. 2012, Yogurtcu and Johnson 2018). The length and amino acid pattern of the IDRs control the sampled space around the membrane and the ability to engage with binding partners (“fly-casting”) (Shoemaker et al. 2000). PTMs on the IDRs regulate membrane-binding (Araya et al. 2022) and modulate signaling cascades (Wright and Dyson 2015). Multivalent low-affinity interactions of membrane-bound IDRs could also drive clustering and phase separation (LLPS) at the membrane (Ditlev 2021). These distinct mechanisms or their blends play a pivotal role in organellar remodeling (Cornish et al. 2020).

Selective ER-phagy, a critical homeostatic pathway, orchestrates the targeted degradation of the endoplasmic reticulum (ER) via autophagy, particularly under stress or ER expansion (Reggiori and Molinari 2022). ER-phagy is emerging as a crucial catabolic pathway that (i) rapidly mobilizes nutrients during starvation (Reggiori and Molinari 2022), (ii) actively regulates the size and shape of the ER (Xu et al. 2021), (iii) ensures clearance of aberrant or aged proteins and lipids (Ferro-Novick et al. 2021), (iv) is co-opted by pathogens to invade host cells (Li et al. 2021), and (v) is deregulated in an increasing number of diseases (Hübner and Dikic 2020). Integral to this process is the recruitment of the phagophore membrane to the ER-phagy sites by distinct membrane-bound receptors (Gubas and Dikic 2022). FAM134B/RETR1, and all known ER-phagy receptors, including SEC62, RTN3, CCPG1, and TEX264, are ER-resident integral membrane proteins that house LIRs in their cytoplasmic IDRs (Reggiori and Molinari 2022). The IDRs serve as flexible and extensible connections between their ER anchor and the LC3/ATG8 proteins covalently attached to the phagophore membrane, which results in the recruitment of the autophagic machinery to the ER.

Selective ER-phagy closely intertwines with ER membrane remodeling. As autophagic membranes are engaged, the ER membrane undergoes large-scale shape changes, leading to fragmentation and engulfment into the autophagosomes (Gubas and Dikic 2022). Recent advances combining cell biology methods, *in vitro* membrane remodeling assays, and extensive MD simulations have identified and mapped the curvature induction, sensing, and sorting functions of the ER-phagy receptors. For instance, FAM134B, RTN3, and TEX264 have membrane-sculpting transmembrane hairpins and amphipathic helices (Reggiori and Molinari 2022). The transmembrane domains of these receptors, particularly the RHDs, adopt a dynamic wedge shape membrane inclusion, sculpting the ER-membrane (Bhaskara et al. 2019). Further, their curvature sensing functions trigger protein sorting and clustering to nucleate membrane buds spontaneously (Siggel et al. 2021). This process is further fine-tuned and accelerated by the ubiquitination of receptors (González et al. 2023) to form large membrane-associated homomeric and heteromeric RHD clusters (Foronda et al. 2023) to enhance ER-phagy. However, the precise functions and role of IDRs within ER-phagy receptors in these various steps remain unclear. The importance of IDRs, especially in the context of membrane remodeling, has not been addressed.

Studying remodeling properties of membrane-associated and membrane-anchored IDRs presents several challenges. The IDR conformations can be variable in solution and membrane-bound states due to differential interactions with solvent and distinct lipid species (Van Der Lee et al. 2014, Cornish et al. 2020, Has et al. 2022). The large conformational entropy of the membrane-bound IDRs is favored by underlying bent and curved bilayer geometry, as demonstrated experimentally for AP180, amphiphysin, and epsin (Zeno et al. 2018, 2019). The increased surface density of IDRs can also perturb bilayers (Stachowiak et al. 2012). IDRs crowded on one side of the membrane surfaces can induce substantial lateral pressure in one of the leaflets (Snead et al. 2017, Houser et al. 2022), resulting in dramatic curvature induction. The interplay between the electrostatic and entropic effects, the presence of other membrane-shaping elements (*e.g*., transmembrane hairpins or amphipathic helices (AHs)), and the crowding effects of tethered IDRs are often challenging to decouple in experiments (Houser et al. 2022). With unmapped relative contributions, the importance of IDRs is often undermined in membrane remodeling.

Here, we aim to decouple the diverse roles of the IDRs in ER-phagy receptors. We use extensive modeling and simulations of the IDR of FAM134B, a well-studied ER-phagy receptor model system, to investigate how the IDR structure and dynamics influence local membrane properties. To this end, we build detailed structural ensembles of the IDR. We use MD simulations to explore its conformational landscape in solution and varied membrane-anchored states. We identify that the FAM134B-IDR adopts alternate conformational states based on the membrane anchor. Our simulations reveal that IDR alone can induce and sense membrane curvature. This effect accentuates the RHD-mediated curvature induction and curvature sensing functions to promote efficient clustering, nucleating membrane buds faster, thereby boosting selective ER-phagy. Our findings suggest that the C-terminal IDR modulates the membrane-shaping functions of the RHD intricately along the various phases of ER-phagy.

## II. RESULTS

### A. Structure of FAM134B-IDR

The well-characterized RHD of FAM134B (Bhaskara et al. 2019) is flanked by two intrinsically disordered fragments on the cytosolic side of the ER membrane, *i.e*., a smaller N-terminal IDR (1–79) and a longer C-terminal IDR (261–497) that houses the functional LIR motif (^453^DDFELL^458^; Fig. 1a). Analysis of the residue composition of these segments revealed several contiguous stretches of 3 to 5 charged residues (red or blue regions; Fig. 1a). We compared the charge characteristics of the C-terminal disordered fragment (referred to hereafter as IDR) of the FAM134 family along with the cytosolic IDRs of other ER-phagy receptor families (Supplementary Fig. S1; Supplementary Table S1). By quantifying the distribution of fractional positive and negative charges within the IDR, we grouped the ER-phagy receptors based on the Das-Pappu classification (Das and Pappu 2013) for IDRs (Fig. 1b). We found that they predominantly correspond to the boundary region (Region 2) between weak and strong polyampholytes, indicative of adopting chimeric structures between more globule-like and partially open tadpole or coil structures (Fig. 1b). Only the SEC62-IDRs correspond to Region 3, indicating more coil-like structures of a strong polyampholyte.

**FIG. 1.**
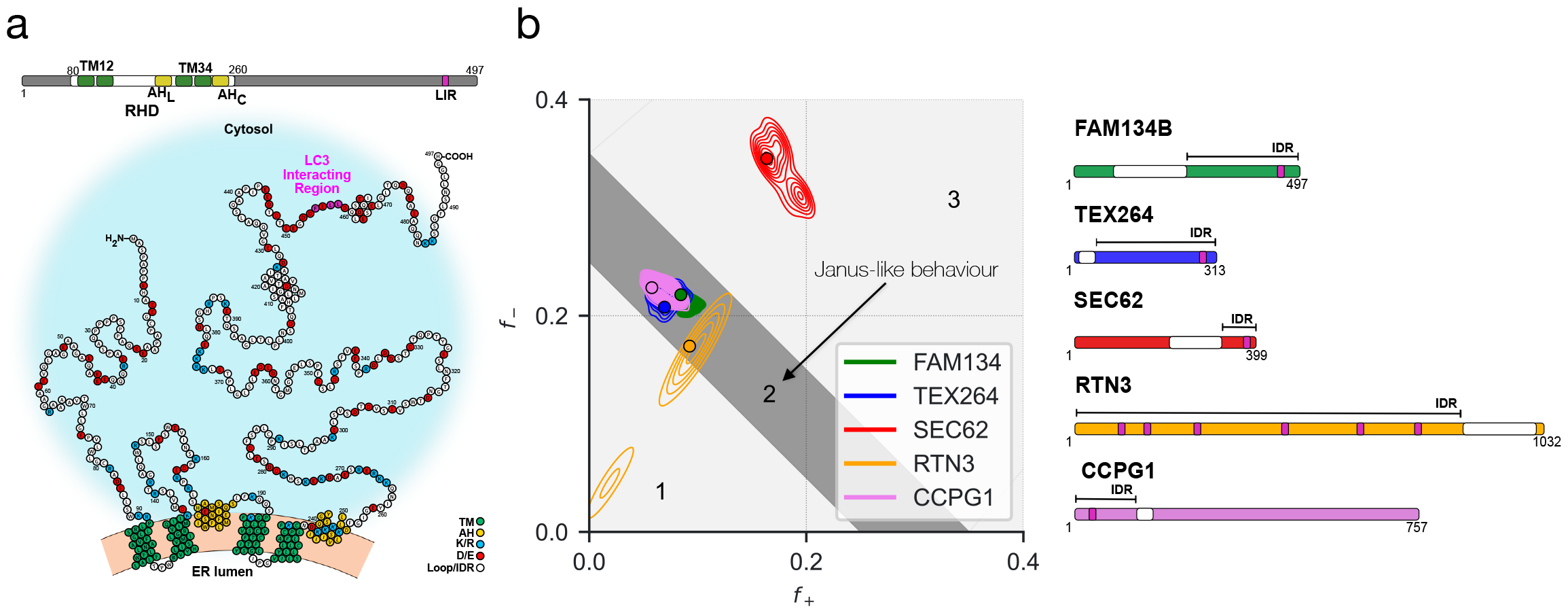
IDRs of selective ER-phagy receptors. (**a**) Domain structure (Top) and topology (Bottom) of full-length FAM134B. The topology diagram shows the organization of the membrane-bound RHD (green and yellow; 80–260) in the curved bilayer (orange). The RHD is flanked by the N-terminal (1–79) and C-terminal (261–497) disordered regions on the cytosolic side of the bilayer. Charged residues (blue or red) and the functional LC3 interacting region (purple) within the IDRs are highlighted. (**b**) Das–Pappu diagram (Das and Pappu 2013) (left) plotting the fraction of positive (*f*_+_; K and R) against the fraction of negative (*f*_−_; D and E) residues for ER-phagy receptors (right). Colored contours show the probability densities estimated from 500 homologous IDR segments corresponding to each known human ER-phagy receptor (filled circles). Regions 1 and 3 of the phase diagram correspond to IDRs with polar tracts (globular or tadpole-like structures) and polyampholytes (coils, hairpins, and chimeric structures), respectively. Region 2 (dark gray) is the transition zone, representing Janus-like quality adopting collapsed or expanded IDR structures.

We analyzed the FAM134B sequence to examine the predicted features of the IDR (Supplementary Fig. S2a). An analysis of the AlphaFold2 (AF) model (Supplementary Figs. S2b–c) of the IDR revealed a predominantly coil-like structure with three distinct high-confidence regions with more local contacts (pLDDT ≥ 80; R1, R2, and R3 in Supplementary Fig. S2b), indicating a partial local residual helical structure for the IDR. Furthermore, these regions also correspond to the predicted PPI sites of FAM134B (MoRFs in Supplementary Fig. S2a), which indicates possible functional constraints to preserving the local helical structure.

Next, we obtained a structural ensemble for the IDR of FAM134B in solution by running 1 μs of atomistic MD simulation starting from the AF model (Fig. 2a; Supplementary Table S2; Supplementary Movie SM1). We found that the IDR ensemble populated relatively compact structures in solution (Fig. 2a; Supplementary Movie SM1) with end-to-end distances of *R*_*e*_ ≈ 4 ± 2 nm and radii of gyration of *R*_*g*_ ≈ 2.6 ± 0.5 nm (mean ± SD). The corresponding 2D free energy surface is shown in Fig. 2b. The local secondary structure remained predominantly coil-like (Fig. 2c, see *P*_C_; blue curve). However, the three residual helical stretches (R1, R2, and R3) remained relatively stable (up to 1 μs), with only the N-terminal R1-helix undergoing partial unfolding (Fig. 2c, see *P*_H_; green curve). We detected that distinct pairwise residue interactions (apolar: S366-L373, L458-L463, and salt bridge: K376-D381) contribute to the stability of the compact structure of the IDR in solution (Supplementary Fig. S2d).

**FIG. 2.**
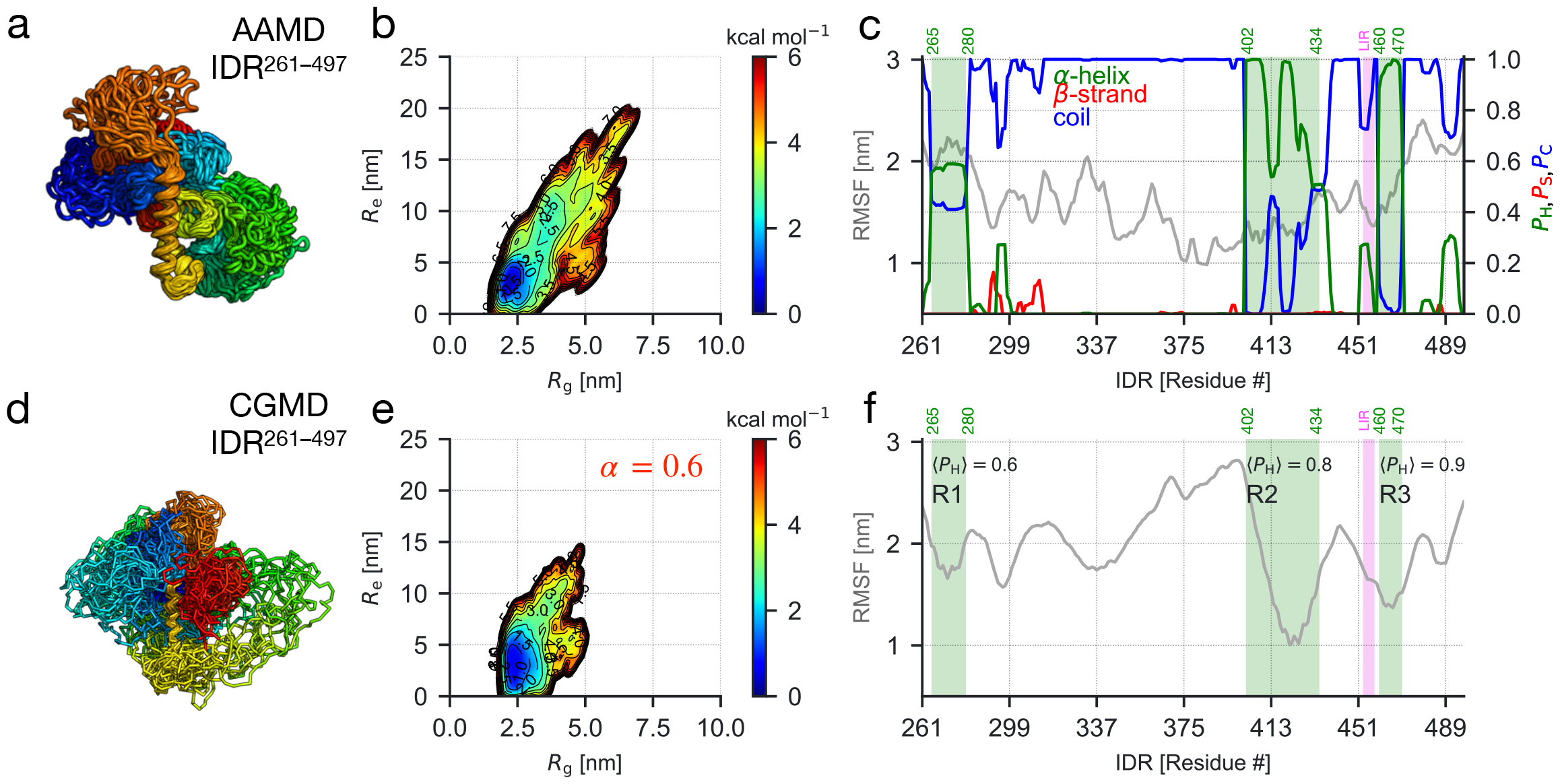
Structural models for FAM134B–IDR in solution. (**a**) IDR ensemble (30 representative structures; rainbow colors) in a dilute solution sampled in atomistic MD simulations. (**b**) Free energy landscape for the IDR ensemble along the radius of gyration (*R*_*g*_) and end-to-end distances (*R*_*e*_) from atomistic MD simulations. (**c**) Root mean square fluctuations (RMSF; grey curve) and secondary structure propensities (helix: green, strand: red, and coil: blue) along the IDR length were obtained from all-atom simulations to quantify the local conformational variability. (**d**) Representative IDR structures from coarse-grained MD simulations obtained after scaling protein-protein interaction strength (*α* = 0.6) to calibrate the (**e**) coarsegrained free energy landscape. (**f**) Regions R1, R2, and R3 (green-shaded) display high local helix propensity (⟨*P*_H_⟩≥0.5) and reduced fluctuations in coarse-grained simulations.

Subsequently, we constructed an effective coarsegrained (CG) model of the IDR to study its motions and its coupling to the RHD. We used the Martini model with reduced protein-protein interaction strength (Figs. 2d–f; Supplementary Table S2; see Methods) to model the IDR. We used alpha-scaling (Stark et al. 2013, Benayad et al. 2020) to vary the PPI strength in a range between *α* = 1.0 (full interaction) to *α* = 0.3 (30% interaction). In this way, we gradually reduced the stickiness of the Martini model, increasingly sampling less compact and more open conformations of the IDR (Supplementary Fig. S3a–d). At *α* = 0.6, we found the free energy surface of the IDR extracted from the scaled coarse-grained MD simulations was consistent with that obtained from atomistic MD simulations (Fig. 2b vs. 2e; Supplementary Fig. S3c; Supplementary Movie SM2). Moreover, at *α* = 0.6, we also found that the values of *R*_*g*_ and *R*_*e*_ averaged over the ensemble also matched closely (Supplementary Figs. S3c, S3e, and S3f). For FUS, similar *α* values gave phase behavior consistent with experiments (Benayad et al. 2020). The good overlap of the freeenergy surfaces of the all-atom and coarse-grained IDR ensembles encouraged us to use *α* = 0.6 to model the effective interactions of the IDR in all subsequent MD simulations.

### B. FAM134B-IDR collapses onto the RHD

To determine if the tethering of the IDR at its N-terminus influences its ensemble properties, we modeled two membrane-anchored IDRs. The N-terminus of the IDR was linked to either KALP_25_, a model transmembrane helical peptide (KALP_25_–IDR), or to the RHD of FAM134B (80–497; RHD–IDR). We then embedded these two constructs in flat model bilayers containing POPC lipids and initiated MD simulations (Supplementary Table S3; Supplementary Movie SM3). The IDR anchored to the KALP_25_ peptide adopted more extended structures (Fig. 3a) with a larger conformational landscape (Fig. 3b), and high flexibility (Fig. 3c). Further, this IDR ensemble displayed high asphericity (≈ 0.4 ± 0.2; Supplementary Fig. S4a), enclosed a much larger hydrodynamic volume above the membrane (grey envelopes in Figs. 3a,d and corresponding radii, *R*_*T*_ ≈ 4.5± 2.0 nm; Supplementary Figs. S4b-c) as opposed to compact conformations of the IDR in solution. By contrast, the IDR tethered to the membrane-bound RHD collapsed quickly (within the first 300 ns) and adopted more compact structures on top of the RHD (Figs. 3d; Supplementary Figs. S4a-c) akin to IDRs in solution. We found that the extent of collapse and compaction of the RHD-anchored IDR, albeit smaller than solution-phase IDR, was consistent in three different replicates, indicating that more specific interactions stabilized this intermediate structure.

**FIG. 3.**
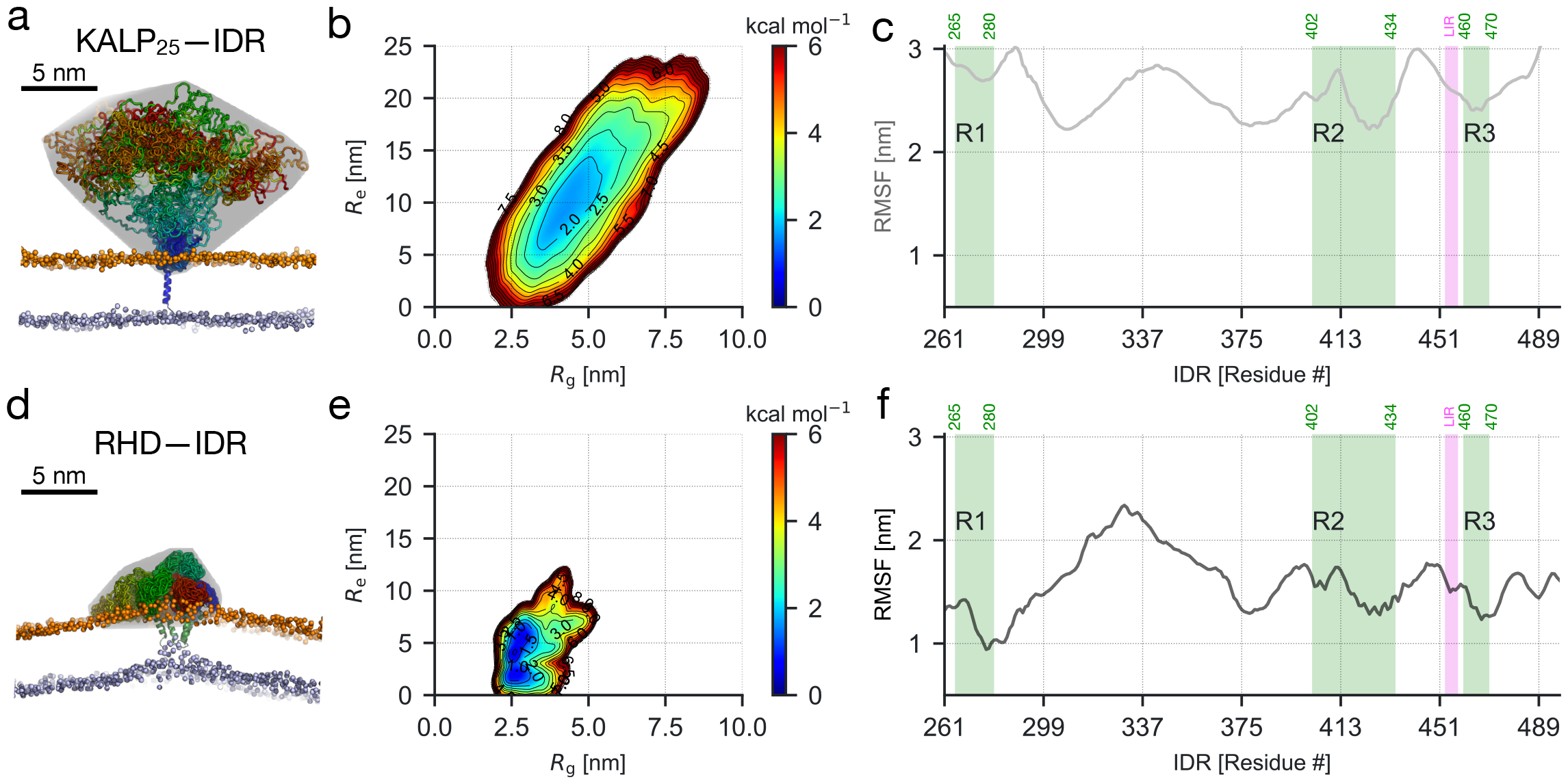
Membrane anchoring influences the conformation of IDRs in coarse-grained MD simulations. Ensemble properties of membrane-anchored IDRs were obtained by connecting them in tandem to the C-terminus of (**a–c**) the KALP_25_ peptide (KALP_25_–IDR) or (**d–f**) the FAM134B-RHD structure (RHD–IDR). (**a, d**) 30 representative conformations (rainbow-colored) of each IDR ensemble aligned to the membrane-bound KALP peptide or RHD. Grey envelope shows their corresponding hydrodynamic volumes (see Supplementary Fig. S4) on the cytosolic side. (**b, e**) Free energy surfaces of IDRs linked to KALP_25_ and RHD along *R*_*g*_ and *R*_*e*_. (**c, f**) KALP_25_–IDR displays larger fluctuations, sampling a larger conformational space. By contrast, the RHD induces compaction of the tethered IDR, inducing the formation of a membrane scaffold stabilized by specific intra-molecular and bilayer interactions of the IDR (see Supplementary Fig. S6).

An analysis of the contact maps of the IDR ensembles averaged from MD simulations (Supplementary Figs. S5a-c) revealed that the overall character remained preserved. Still, the intensities of specific residue-wise contacts varied under the different contexts of the IDR studied (unbound and anchored). We found that the compaction of the IDR in solution and partially in RHD-anchored IDR are predominantly due to the increased contacts of R2 and R3, regions that display residual helical structure. Further, we quantified RHD–IDR interactions driving the compaction and stabilization of the tethered IDR (Supplementary Fig S6; Supplementary Movie SM4). We found that the residues 306–351 of the IDR consistently displayed contacts with the linker region of the RHD containing an amphipathic helix (AH_L_; see Supplementary Fig. S6a), resulting in IDR compaction. The collapse of the IDR was slowed but overall preserved when we altered the interaction strength of the IDR (using *α* = 0.1; Supplementary Fig. S6b) or changed R1, R2, and R3 from helix to coil conformation (Supplementary Fig. S6c). Additionally, we found that transient interactions of hydrophobic residues of the IDR with the membrane lipids (Supplementary Fig. S6, right) contributed to the overall compact state of the IDR. Two short hydrophobic stretches (280–300 and 390–400) interacted consistently with POPC lipids, causing the IDR to scaffold over the RHD (Supplementary Fig. S6a, right).

### C. FAM134B-IDR induces membrane curvature

To determine if the structure and dynamics of membrane-anchored IDRs influence the local membrane shape, we monitored the local height of the bilayer, *h*(*x, y*), and its associated curvature field, *H(x, y)*, under the influence of periodic boundary conditions (see Methods). We used membrane containing KALP_25_ and RHD without anchored IDRs as negative and positive controls, respectively. As expected, the bilayer with embedded KALP_25_ remained flat with minimal height fluctuations (Fig. 4a). However, when linked to the IDR (KALP_25_–IDR), we found a slight bump in the height profile of the membrane consistent with the induction of a small positive membrane curvature (Fig. 4b). This indicated that the IDR could induce positive membrane curvature locally. We also confirmed our previous computations (Bhaskara et al. 2019) that the RHD alone perturbed the bilayer by causing a local bulge with strong positive curvature (Fig. 4c). The membrane-shaping effect was greatly amplified upon tethering the IDR to the RHD, resulting in a distinct membrane bulge (Fig. 4d).

**FIG. 4.**
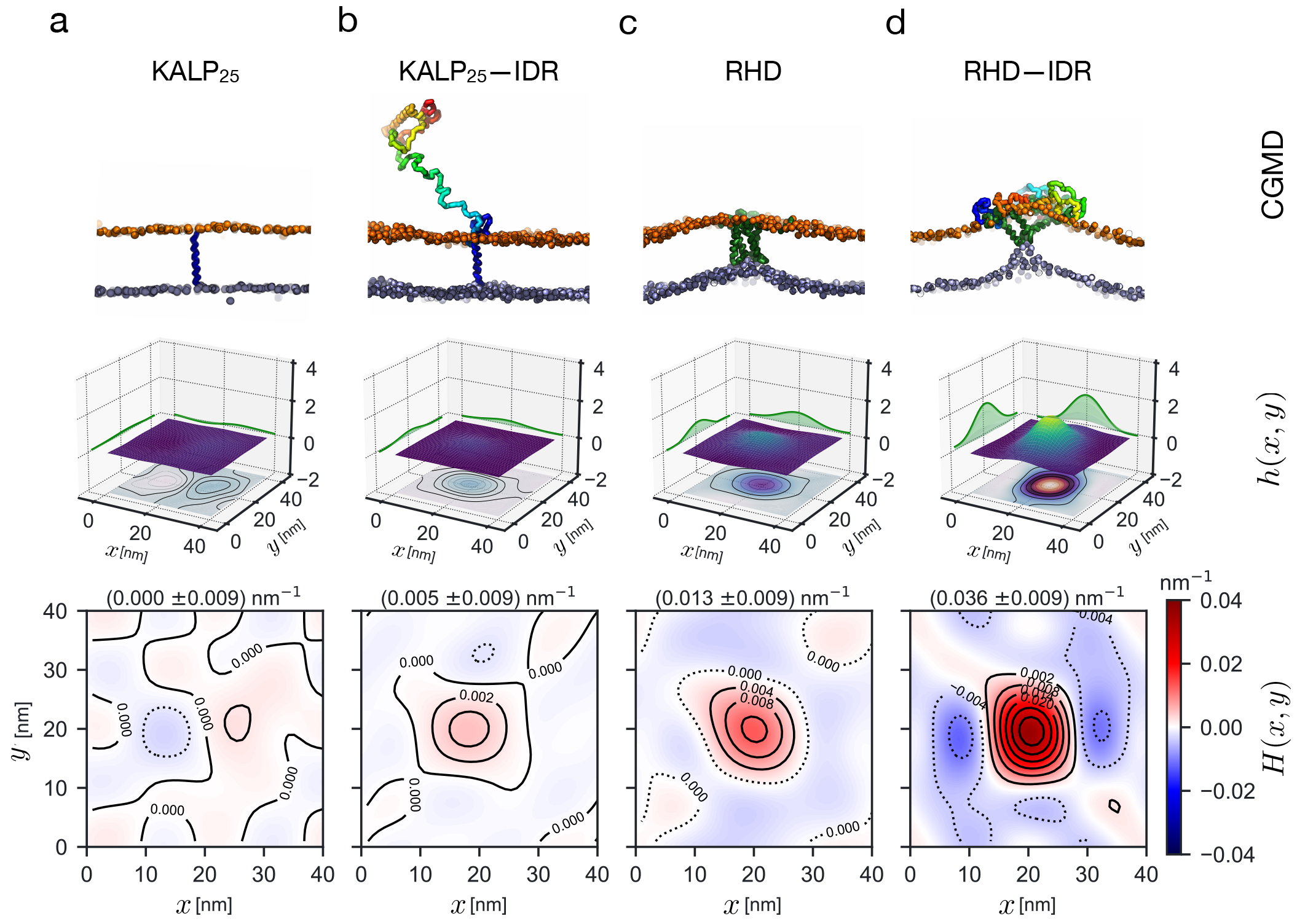
Membrane-anchored IDRs induce membrane curvature in coarse-grained MD simulations. Square bilayer patches (38 × 38 nm^2^) under periodic boundary conditions containing (**a**) KALP_25_ (negative control), (**b**) KALP_25_–IDR (**c**) FAM134B–RHD (positive control) and, (**d**) RHD–IDR perturb bilayer structure differently. (**Top**) Side view snapshots at 5 *μ*s showing the characteristic local membrane shape represented by lipid head groups from both leaflets (orange and blue PO4 beads) around the membrane inclusion. (**Middle**) Local membrane shape is approximated by a height function *h*(*x, y*) of the midplane (height contour map along *xy* plane). Projections onto the *xz* and *yz* planes show the shape profile around the inclusion. (**Bottom**) Contour maps of the averaged curvature profiles *H*(*x, y*) for each system show the extent of curvature induction around the inclusion centered in the box. The maximum value of curvature fields ⟨*H*_max ⟩_ and ± SD is shown at the top. Midplane *h*(*x, y*) and corresponding *H*(*x, y*) are computed by fitting 500 individual frames at 2-ns intervals over the last 1 *μ*s of representative trajectories.

To further test the role of IDRs in inducing membrane curvature, we lowered the barrier for local bulging by increasing the bilayer asymmetry (Δ*N* = *N*_*upper*_ *N*_*lower*_ = [100, 300, 500]; Supplementary Fig S7). We found that the RHD–IDR-induced membrane curvature increased with the extent of bilayer asymmetry. This pattern was also observed for RHD alone but moderately. By contrast, the presence of other proteins did not show a steady increase in the bilayer curvature with increasing bilayer asymmetry (Supplementary Fig. S7).

We further elucidated the role of the IDR in direct membrane remodeling by re-analyzing previous experimental results from FAM134B-induced *in vitro* liposome remodeling (Bhaskara et al. 2019, González et al. 2023). In these experiments, we had reconstituted empty liposomes (*d* ≈ 200 nm) with purified proteins (GST-tagged FAM134B–RHDs, full-length FAM134B-WT, and FAM134B–17KR mutant, where all 17 RHD sites have K → R mutations) and imaged them by negative-stain transmission electron microscopy (nsTEM; Supplementary Fig. S8). We found that the FAM134B–WT and the FAM134B–17KR mutant with intact IDRs drastically remodeled larger liposomes into smaller vesicles (*d* = 28±16 nm and *d* = 27±8 nm). By comparison, the RHD alone (Supplementary Fig. S8) retained liposome remodeling behavior (*d* = 57±25 nm), albeit less than RHDs flanked by IDRs. Our simulations on RHD–IDR are consistent with these observations and explain the difference in the observed size distribution of proteoliposomes.

### D. FAM134B-IDR senses membrane curvature

Next, we determined the intrinsic curvature preference of proteins with and without tethered IDRs by simulating a buckled bilayer under tension (see Methods; Supplementary Table S4; Supplementary Movie SM5). The membrane buckle presents a sinusoidal carpet-like folded structure with a range of local mean curvatures (*H*(*x, y*)= − 0.07 to +0.07 nm^−1^) under periodic boundary conditions. By lateral diffusion (Bhaskara et al. 2019) or dissociation and re-association (Rao et al. 2024), membrane-bound proteins preferentially sample different local curvature environments, allowing us to directly quantify their curvature preference (Fig. 5).

**FIG. 5.**
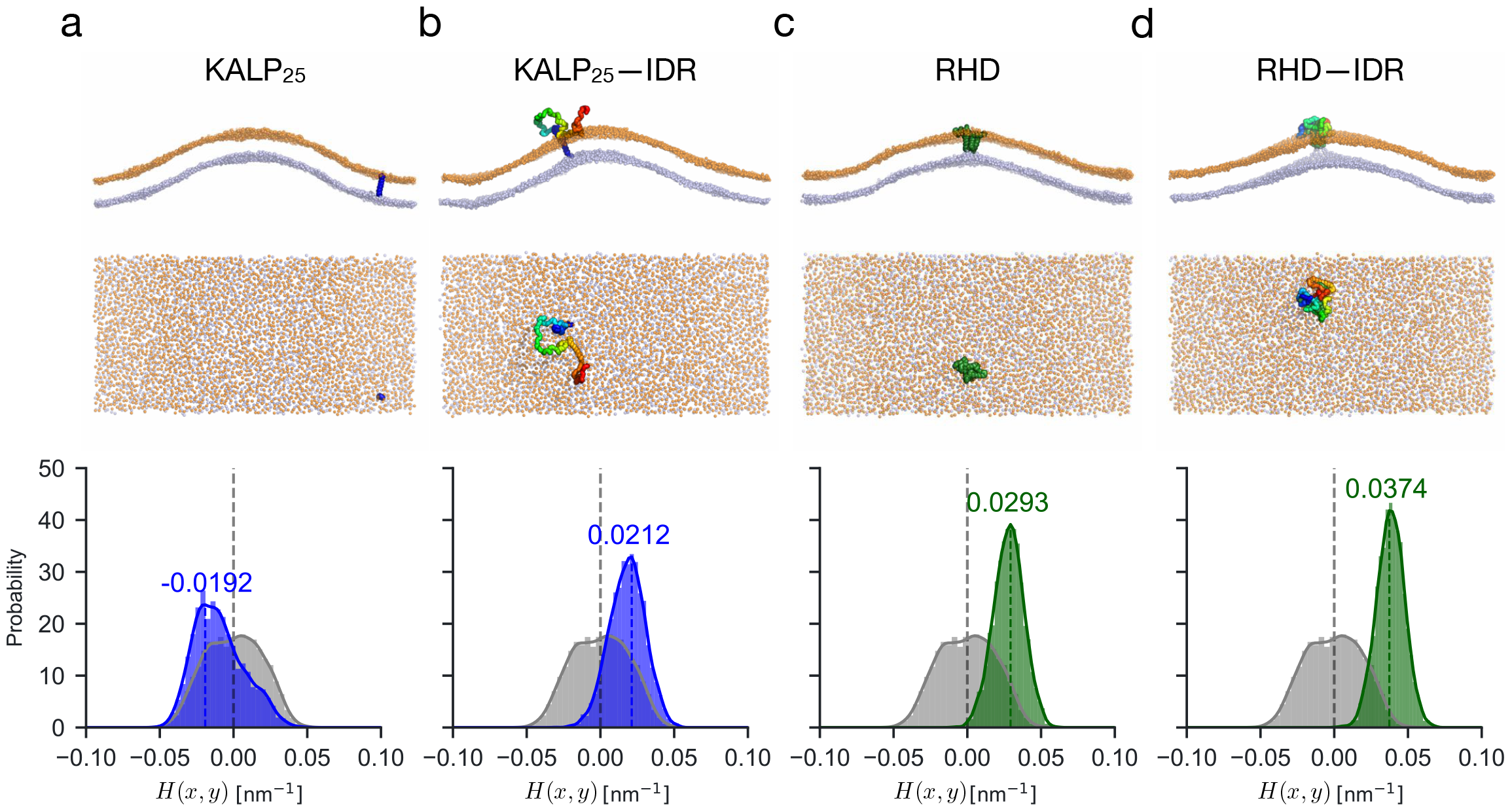
Membrane-anchored IDRs sense membrane curvature. Rectangular membrane buckles (58 × 27 nm^2^) were embedded with proteins (**a**) KALP_25_ (negative control), (**b**) KALP_25_–IDR (**c**) FAM134B–RHD (positive control) and, (**d**) RHD–IDR were embedded in rectangular membrane buckles (60 ×30 nm^2^) and subjected to coarse-grained MD simulations to quantify their curvature preferences. (**Top**) Snapshots (side and top views) at the end of the 10 *μ*s runs show the preferred position of the protein along the buckle. (**Bottom**) Histograms of the mean curvature *H*(*x, y*) sampled by the center-of-mass positions of the proteins (transmembrane regions only, KALP_25_ in blue; RHD in green) indicate curvature preference along the buckle. The IDR-containing proteins sample regions of high mean curvature and preferentially occupy the top of the buckle. For reference, the distribution of local mean curvature values on the empty buckled membrane (grey) was estimated by random sampling of points in the *xy* plane, ignoring small curvature corrections. The time-averaged values of *H*(*x, y*) are also highlighted.

In control simulations with the KALP_25_, we observed a slight preference for surfaces with negative mean curvature (*H*(*x, y*)= − 0.019 nm^−1^; Fig. 5a) akin to structures at the bottom of the buckle with negative Gaussian curvature (*K*_*G*_ ≤ 0; Supplementary Fig. S9a). Consistent with previous computations, we found that the RHD alone strongly preferred regions of high local mean curvature (*H*(*x, y*) = 0.029 nm^−1^; Fig. 5c; Supplementary Fig. S9c) occupying the top of the membrane buckle (*K*_*G*_ > 0 corresponding to ellipsoidal vesicle shapes). Interestingly, by tethering an IDR to the KALP_25_ peptide, we found that the curvature preference of the molecule changed drastically. The KALP_25_–IDR displayed biased diffusion and migrated towards the top of the buckle with a preference for regions with high positive mean curvature (*H*(*x, y*) = 0.021 nm^−1^; Fig. 5b) akin to the tubular geometry on top of the buckle (*K*_*G*_ ≥ 0; Supplementary Fig. S9b). Upon tethering the IDR to the RHD, we found that the curvature profile of the buckle altered more swiftly, causing an enhanced preference for positively curved surfaces on top of the buckle (*H*(*x, y*)= 0.037 nm^−1^; Fig. 5d). This enhanced curvature preference exceeds the RHD’s curvature preference, is bi-directional (*K*_*G*_ *≫* 0; Supplementary Fig. S9d), and further deforms the top of the buckle.

### E. FAM134B-IDR boosts RHD-mediated membrane budding

Motivated by the above findings, we investigated the impact of membrane-anchored IDRs on RHD-induced budding. Previously, we probed the effect of RHD clustering on membrane shape (Siggel et al. 2021). As in this earlier study, we modulated the kinetic barrier and energetic driving force for forming highly curved bud shapes by varying the bilayer leaflet asymmetry and protein concentration. We initiated simulations from flat metastable asymmetric bilayers (2–13% leaflet asymmetry) containing different numbers *n*_Prot_ = [3, 6, 9] of KALP_25_ peptides, KALP_25_–IDRs, RHDs, or RHD–IDR molecules (see Methods; Supplementary Fig. S10; Table S5). Tracking individual proteins (z-position) and their interactions (clusters) along with the box width (*L*_*x*_) in these simulations provided an excellent set of order parameters to characterize protein clustering, membrane shape changes, and other features associated with spontaneous budding (Fig. 6).

**FIG. 6.**
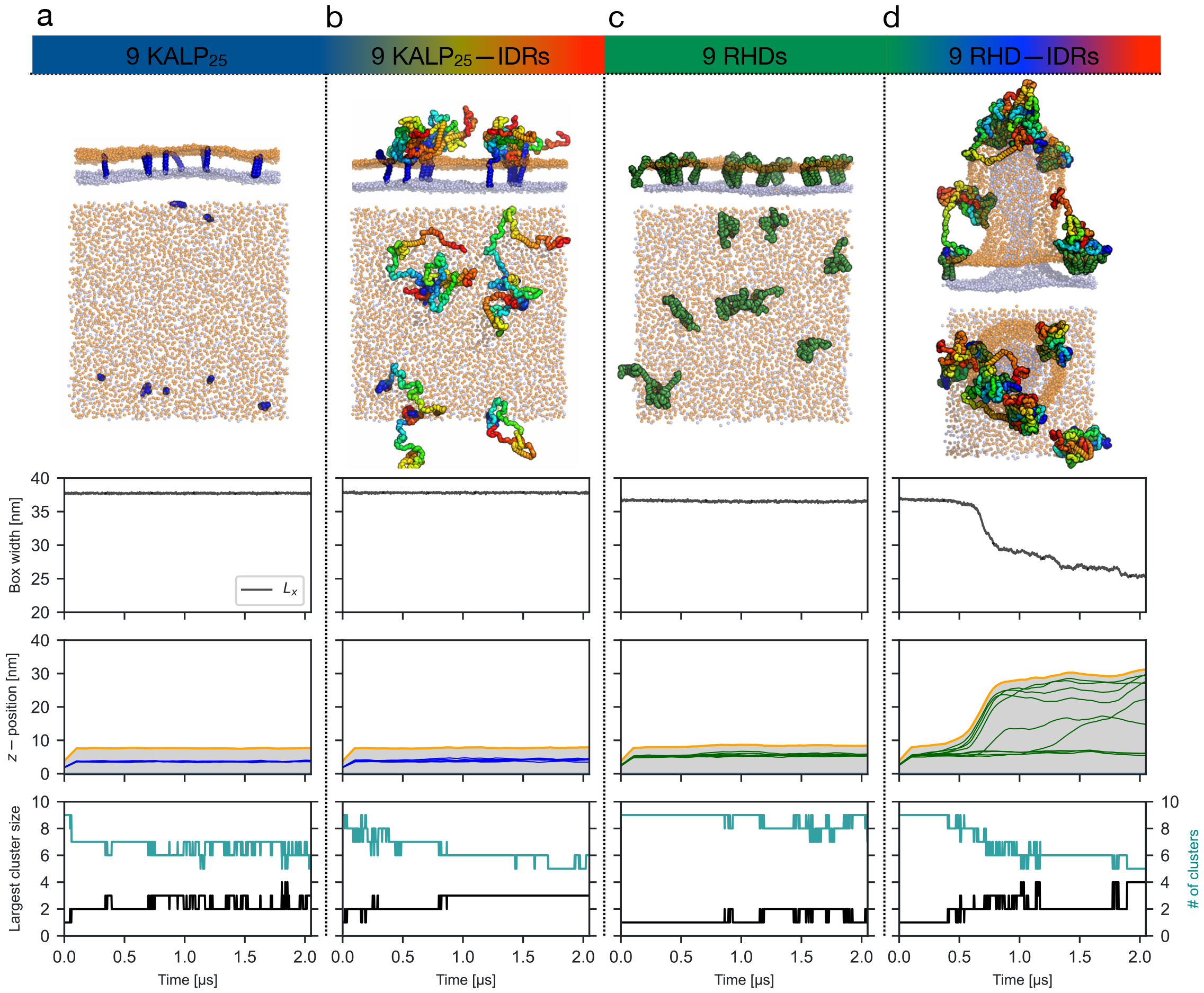
Membrane-anchored IDRs boost membrane budding. In coarse-grained MD simulations, we embed flat square asymmetric bilayer patches (Δ*N* = 500; 38 × 38 nm^2^) with nine copies of (**a**) KALP_25_ (negative control), (**b**) KALP_25_–IDR (**c**) FAM134B-RHD (positive control), and (**d**) RHD–IDR. (**Top**) Snapshots showing side and top views of the organization of proteins at the end of 2 *μ*s runs. (**Middle**) The central two rows display the time traces of order parameters informing on spontaneous membrane budding events. Sharp drops in the box width (*L*_*x*_) and rapid vertical displacement *z* of individual proteins (COM positions of transmembrane segments; blue and green lines), along with a rise in the highest points in the bilayer (orange lines) and the intervening range (light grey shaded region), indicate membrane budding. (**Bottom**) The number of segregated protein clusters and the size of the largest protein cluster quantify the extent of clustering, determining the kinetics of budding. IDRs mediate inter-protein interactions and enhance RHD–IDR clustering to induce spontaneous membrane buds faster. We note that on a longer time scale, the RHDs alone also induce membrane budding (see Supplementary Figs. S10–11).

We found that bilayers containing KALP_25_ peptides or KALP_25_–IDRs did not induce membrane budding under all simulation conditions (Figs. 6a–b; Supplementary Figs. S10a–b; Supplementary Table S5; Supplementary Movie SM6). Consistent with our previous studies, RHDs, and RHD–IDRs induce spontaneous membrane buds by clustering (Figs. 6c–d; Supplementary Movie SM6). The number of budding events within the three replicates increased with protein concentration and bilayer asymmetry (*n*_Prot_ and Δ*N*; Supplementary Figs. S10c–d; Supplementary Table S6). For RHDs alone in the absence of IDRs, we found infrequent budding that occurred at a much slower rate (*N*_buds_ = 1/3; *K*_RHD_ = 20 ns^−1^; Supplementary Fig. S11a). By contrast, the RHDs with tethered IDRs induced membrane buds more frequently at faster rates on the MD timescale (*N*_buds_ = 3/3; *K*_RHD–IDR_ = 57 ns^−1^; Supplementary Fig. S11b). Further, nine RHDs were required to induce budding at intermediate leaflet asymmetry (6.5–7.5%; Δ*N* = 300; Supplementary Fig. S11a; Supplementary Movie SM6).

By contrast, under the same conditions, six RHD–IDRs were sufficient to induce budding repeatedly (Supplementary Fig. S11b). We express the driving force for membrane budding by measuring the rate from observed waiting times for bud formation (Supplementary Table S6). We estimate that the IDRs of FAM134B accelerated the kinetics of RHD-mediated spontaneous budding by a factor of 3.3 and 2.0 for bilayers with intermediate (Δ*N* = 300) and high (Δ*N* = 500) asymmetry, respectively. In simulations containing multiple copies of RHD–IDRs, we found that IDR dynamics increased inter-protein contacts in the solution phase (Supplementary Fig. S12a). Mapping IDR–IDR interactions from MD simulations identified more frequent contacts of residual helical structures R2 and R3. These contacts promoted RHD-clustering, leading to accelerated kinetics of budding.

At high asymmetry (Δ*N* = 500), bilayers containing KALP_25_s or KALP_25_–IDRs remained nearly flat, indicating that the KALP_25_ peptide clusters within the bilayer were insufficient to overcome the barrier for budding (Supplementary Figs. S10a–b). Although KALP_25_–IDRs did not induce spontaneous buds, the solution-phase IDR–IDR interactions promoted faster protein-clustering (Supplementary Fig. S12b), resulting in local membrane bulges with marginal positive curvatures (Supplementary Fig. S13). To test if such structures could drive budding ultimately, we increased the protein surface density from 6233 molecules per μm^2^ to 13850 molecules per μm^2^. We applied slight positive lateral pressure (*P*_*xy*_ =2 bar), lowering the barrier for budding (Supplementary Fig. S14). Despite these favorable conditions, the bilayer with KALP_25_ peptides remained flat and resisted buckling. By contrast, the IDR–IDR interactions of KALP_25_–IDRs broke this metastability by inducing a bulge that then transitioned quickly into a membrane bud, even at low and intermediate leaflet asymmetries (2–7.5%; Δ*N* = [100, 300]).

## III. DISCUSSION

IDRs are ubiquitous in autophagy. Their prevalence in effector proteins during autophagy initiation, autophagosome nucleation, expansion, and maturation provides functional plasticity to distinct molecular processes (Mei et al. 2014). The abundance of IDRs in established and newly identified selective autophagy receptors, coupled with the existence of LIRs, PTMs, and diverse interaction partners, underscores their central role in autophagy (Cristiani et al. 2023). Further, the cytosolic IDRs of membrane receptors exploit their unstructured conformational ensemble, propensity for PTMs, and alternate motif-based binding modes to exert direct control on organellar homeostasis (Van Der Lee et al. 2014).

Modeling IDR ensembles is challenging. Their structural and functional characterization relies heavily on recent advances in experiment (Naudi-Fabra et al. 2021) and theory (Tesei et al. 2024, Lotthammer et al. 2024). In particular, particle-based MD simulations using coarse-grained models offer a powerful means, enabling direct comparison with experimental observables (Shrestha et al. 2021). Despite its limitations (Alessandri et al. 2019), the Martini force field has proven to be highly efficient for simulating protein-membrane systems (Bhaskara et al. 2019, Monticelli et al. 2008, Marrink et al. 2007). However, for several multi-domain proteins and IDRs, it might lead to an overestimation of protein-protein interactions (Stark et al. 2013), resulting in unrealistically compacted regions (Larsen et al. 2020). Current solutions to overcome the stickiness rely on the improvement of interaction models (Zavadlav et al. 2015, Borges-Araújo et al. 2021) and enhanced sampling (Wassenaar et al. 2015).

Scaling of the protein-water (Thomasen et al. 2022) or protein-protein interactions (Benayad et al. 2020) in the Martini model is required to simulate the correct ensemble properties of IDRs, including correct folding (Poma et al. 2017), formation of RNA–IDR complexes (Martin et al. 2021), and the formation of phase-separated condensates by FUS (Benayad et al. 2020). Here, we modeled the IDR of FAM134B using extensive parametrization. By adopting a scaled Martini model optimized against atomistic MD simulations, we provide a refined depiction of the FAM134B–IDR ensemble in solution. By employing hybrid scaling within the same polypeptide, we provide an even more optimized description of membrane-anchored IDRs and their interactions under varied molecular contexts (*e.g*., tethered to different membrane proteins). This allowed us to exploit the efficiency of the Martini model to study the influence of the IDR on the membrane remodeling capacity of ER-phagy receptors (Bhaskara et al. 2019, Siggel et al. 2021, González et al. 2023).

IDR conformations sampled in the coarse-grained simulations (*α* = 0.6) show a quasi-qualitative overlap with all-atom simulations reproducing its conformational landscape. The overall structure and organization of the IDR ensemble, its residual secondary structure, and tertiary interactions are consistent with sequence-based predictions of IDR characteristics (*e.g*., predicted MoRFs, pLDDTs, and binding sites). Anchoring the IDR to bilayers resulted in a context-dependent sampling of conformations. Consistent with its location in the Das-Pappu phase diagram (Das and Pappu 2013), the membrane-anchored IDRs display both expanded (KALP_25_–IDR) and compact states (RHD–IDR), highlighting its context-dependent behavior. The preservation of this feature across the majority of ER-phagy receptors and their homologs strongly suggests a significant functional or regulatory role for the IDRs.

The combination of protein disorder and membrane dynamics is central to signaling (Cornish et al. 2020). Membrane-anchored IDRs mediate signal transduction across bilayers, scaffold signaling complexes, and regulate vesicle trafficking, functions central to ER-phagy receptors (Van Der Lee et al. 2014). Our simulations of IDRs anchored to the bilayer perturb local membrane shape, clearly demonstrating IDR-mediated active membrane curvature induction and sensing. We found that IDRs anchored to membranes adopt extended conformations, increasing their hydrodynamic volume (Stachowiak et al. 2012). Tethering them to flat bilayers also limits the number of accessible conformations, inducing positive membrane curvature through entropic forces (Zeno et al. 2018, Yu and Sukenik 2023) as demonstrated for IDRs in epsin, AP180, and amphiphysin (Zeno et al. 2018, 2019). Alternatively, IDRs with a negative net charge could be electrostatically repelled by a bilayer containing anionic lipids to induce local membrane bending, reducing overall steric hindrance (Zeno et al. 2019). Phosphorylation of the FAM134B–IDR could enhance this effect.

Further, at high concentrations, IDRs induce increased lateral pressure, enhancing membrane deformations (Snead et al. 2016). Steric crowding of IDRs in amphiphysin and epsin induces the formation of buds or tubules (Zeno et al. 2018, Stachowiak et al. 2010). In light of new simulations of the RHD with the C-terminal IDR, we revisited previous *in vitro* liposome remodeling assays (Bhaskara et al. 2019, González et al. 2023) and found that full-length FAM134B-WT and the FAM134B-17KR mutant with preserved C-terminal IDRs are much more efficient in remodeling liposomes (≈ 56% smaller liposomes), highlighting the importance of IDRs in accentuating RHD-mediated curvatures. Consistent with these observations, we found that FAM134B–IDR mediated protein-protein interactions, enhancing the kinetics of receptor clustering and membrane budding. Often, the entropic, electrostatic, and crowding mechanisms collectively influence the structural state of the IDR, thereby modulating local membrane curvature and large-scale remodeling (Houser et al. 2022).

FAM134B–IDR plays a pivotal role in magnifying membrane remodeling during ER-phagy (Fig. 7). The IDR conformations and emergent ensemble properties are directly influenced by tethered RHDs (Fig. 7a). Further, phosphorylation and ubiquitination of specific residues could aid in reversible switching between expanded and compact IDR conformations (Fig. 7b). The entropic force generated by the surface tethering is sequence-encoded and fine-tuned by the immediate environment. The interplay of variable conformational entropy, direct membrane interactions, and RHD scaffolding amplifies curvature induction and sensing with significant implications across various steps of selective ER-phagy (Fig. 7c). The LIR motif, located within the conformationally variable IDR, recruits hATG8 presumably through fly-casting mechanisms. Fuzzy binding modes of multiple LIR–LDS interactions heighten avidity, stabilizing molecular bridges across the ER and the phagophore membrane. Additionally, the Janus-like behavior of the IDRs, along with IDR-IDR interactions, intensifies volume exclusion effects, increasing the effective receptor concentration and expediting membrane budding (Fig. 7d).

**FIG. 7.**
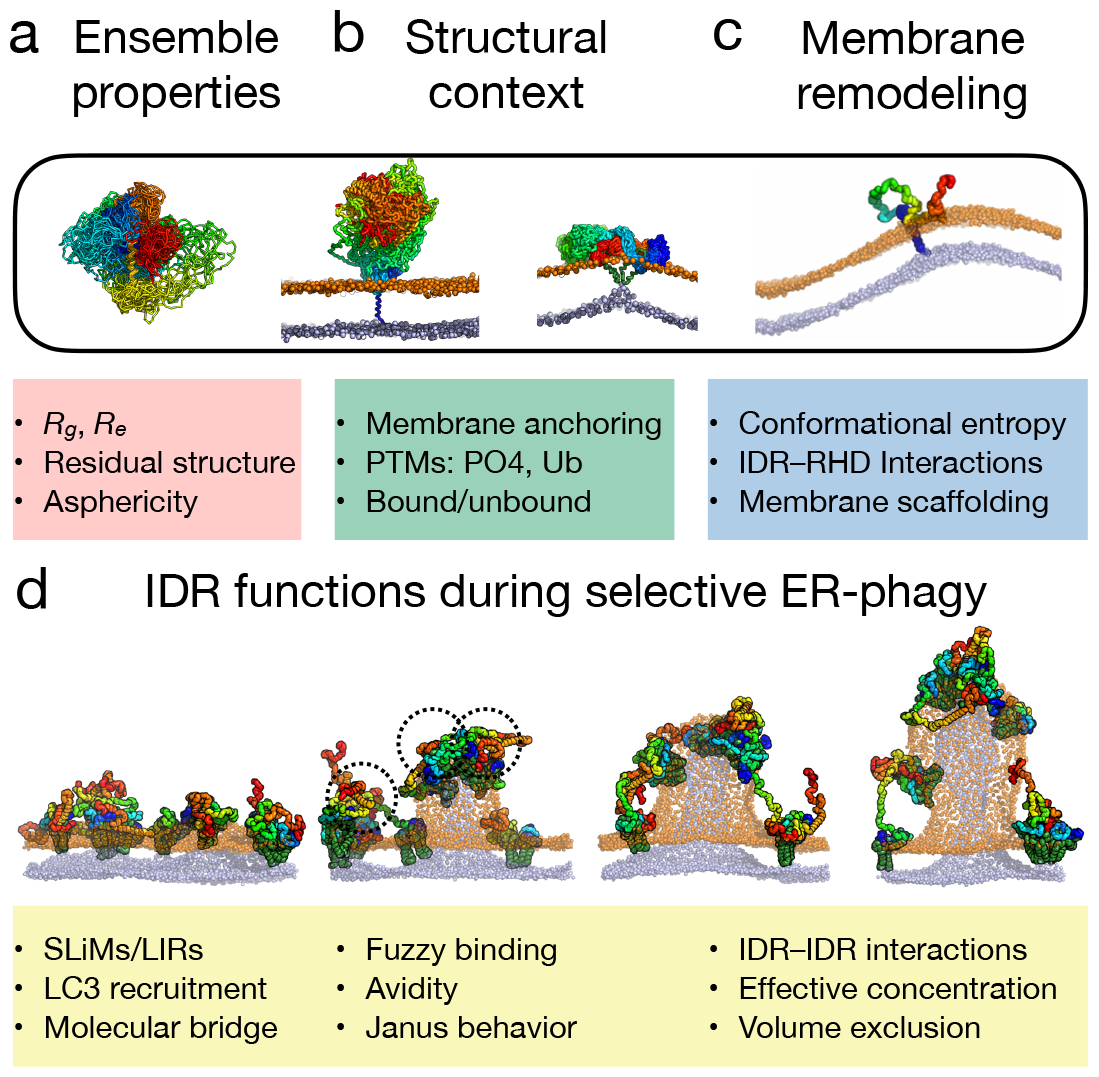
IDRs amplify membrane remodeling during ER-phagy. (**a**) Ensemble properties of the FAM134B–IDR are determined not only by its unique sequence features but are also strongly influenced by the diverse (**b**) molecular, structural contexts and various cellular physiological states. (**c**) IDRs anchored to the RHDs of individual receptors, driven by conformational entropy, can actively induce and sense local bilayer curvature, enhancing the RHD functions. (**d**) The Janus-like behavior of the IDR affects conformational sampling and could influence the fuzzy LIR–LDS binding to hATG8 proteins to enhance avidity during phagophore recruitment. Further, solution-phase IDR–IDR interactions increase effective RHD concentration to drive faster clustering and budding, amplifying large-scale membrane remodeling processes during ER-phagy.

In conclusion, we identified a new role for the IDRs in ER-phagy receptors. We demonstrated how context-dependent ensemble properties of the IDR influence local membrane properties to induce and sense positive membrane curvature actively. Coupled with established RHD functions (Bhaskara et al. 2019, Siggel et al. 2021), the IDRs enhance receptor clustering and hasten membrane budding, thereby augmenting selective ER-phagy. It remains to be seen how multiple regulatory mechanisms coupled to IDR conformational dynamics are actively exploited under varied physiological and stress conditions to modulate overall autophagic flux.

## IV. METHODS

### A. Sequence analysis of IDRs in ER-phagy receptors

IDR segments of well-known ER-phagy receptors: FAM134B, TEX264, SEC62, RTN3, and CCPG1 (Supplementary Table S1) were annotated using D^2^P^2^ (Oates et al. 2012), a consensus predictor combining nine different IDR prediction algorithms (https://d2p2.pro/; Supplementary Figs. S1, S2a). Homologs for all ER-phagy receptors were obtained using psi-BLAST (Altschul et al. 1997) searches against the NR database. Hits (*n* = 500 each) were filtered with an E-value cut-off of 10^−4^, sequence identity range of 30–90%, and a query coverage of ≥ 70%. IDR annotations for the homologs were then mapped from query-hit alignments to compute the fraction of positive (*f*_+_; K and R), negative (*f*_−_; D and E), total charged residues (FCR = |*f*_+_ + *f*_−_), and the net charged sites (NCPR = |*f*_+_ − *f*_−_|). Probability distributions of charge properties of IDR were mapped onto the Das-Pappu plot to identify the most plausible conformational state (*i.e*., polar tracts, polyampholytes, or polyelectrolytes) (Das and Pappu 2013).

### B. Structural model for FAM134B–IDR

Previously, the structure of FAM134B obtained from extensive modeling and simulations was limited to the membrane-bound RHD (Bhaskara et al. 2019). We used AlphaFold2 (https://alphafold.ebi.ac.uk/) to obtain an initial structural model for the C–terminal IDR (residues 261–497) of FAM134B (Jumper et al. 2021). The local per-residue confidence score (pLDDT: predicted Local Distance Difference Test) and PAE (Predicted Alignment Error) were used to assess any residual secondary structures (*e.g*., helical stretches with pLDDT ≥ 50%) and tertiary contacts along the IDR and compared with sequence-based predictions of molecular recognition features from D^2^P^2^ (Supplementary Fig. S2). We connected the IDR structure to the C-terminus of KALP_25_ peptide and RHD (residue 80–260) to model membrane-anchored molecules, KALP_25_–IDR and RHD–IDR, respectively. These models were then energy minimized to remove steric clashes and equilibrated under appropriate solvent conditions.

### C. Molecular dynamics simulations

All MD simulations, including energy minimization, equilibrations (NVT and NPT), and production runs, were performed with GROMACS v2021.5 (Kutzner et al. 2022). A summary of all the MD simulations performed in this study is provided in Supplementary Tables S2– S5. MD simulations of various protein systems were performed in solution (150 mM NaCl), embedded in flat bilayers (symmetric and asymmetric), and embedded in buckled bilayers to study the IDR conformational dynamics, IDR-induced curvature induction, sensing, and large-scale membrane remodeling.

The AF model of the IDR of FAM134B was used as an initial structure to obtain the IDR conformational ensemble in the solution. The IDR in dilute solution was modeled by first centering the IDR in a large periodic dodecahedron box with a distance of 4 nm between the protein and the box edge and then solvating it with TIP3P water containing 150 mM NaCl to allow extensive structural rearrangements. We used the CHARMM36m force field (Huang et al. 2017) for atomistic MD simulations. Electrostatic interactions were modeled with particle mesh Ewald summation and a real-space cutoff of 1.2 nm (Essmann et al. 1995). The LINCS algorithm was used to constrain covalent bonds between hydrogen and heavy atoms (Hess et al. 1997). The Verlet cutoff scheme and the force-switch modifier were used for van der Waals forces with a cutoff of 1.2 nm. The system was first energy-minimized using the steepest descent algorithm and then equilibrated with position restraints on backbone heavy atoms (F = 1000 kJ mol^−1^ nm^−2^) under NPT conditions of constant pressure and temperature, respectively. The system temperature was maintained at 310 K by a Nosé-Hoover thermostat (Nosé 1984a,b, Hoover 1985) with a coupling constant τ_T_ = 1.0 ps. The system pressure was maintained at 1 bar using the isotropic Parrinello-Rahman barostat (Parrinello and Rahman 1981) with a coupling constant of τ_P_ = 1.0 ps, and a compressibility of 4.5 ×10^−5^ bar^−1^. Production runs using a 2 fs timestep for 1 μs were then initiated for 2 replicates with different initial velocities to assess the IDR conformational dynamics.

We used the Martini 2.2 model (Marrink et al. 2007) for coarse-grained molecular dynamics (CGMD) simulations. Four different protein-membrane systems were built and simulated in flat and buckled bilayers *viz*., (i) KALP_25_, a transmembrane helical peptide spanning the bilayer (negative control), (ii) KALP_25_–IDR, where the N-terminus of the IDR is tethered to the KALP_25_, (iii) RHD of FAM134B (positive control) (Bhaskara et al. 2019), and (iv) RHD–IDR, where the N–terminus of the IDR is linked to the C-terminus of the RHD. Initial all-atom models were first assembled for these four protein systems from individual fragment structures (*i.e*., KALP_25_, RHD, and IDR) and then converted to coarsegrained representation using the `martinize.py` script (v2.6) (Monticelli et al. 2008). The local secondary structure of the RHD and the KALP_25_ peptide was preserved by adding backbone constraints obtained from DSSP assignments (Kabsch and Sander 1983). The CG models were then embedded in model bilayers made of POPC (16:0-18:1 PC) lipids using `insane.py` (Wassenaar et al. 2015), followed by solvation using CG water along with 150 mM NaCl.

In all coarse-grained MD simulations, long-range electrostatic interactions were treated using a reaction field with a Coulomb cutoff of 1.1 nm and a dielectric constant ϵ_rf_ of 15. Van der Waals interactions were modeled with a cutoff of 1.1 nm, using the Verlet cutoff scheme and the potential-shift-Verlet modifier. Two rounds of steepest descent energy minimizations (3000 steps each) were performed, first using a softcore potential, followed by an energy minimization without any position restraints. Systems were then equilibrated under NVT conditions with the velocity-rescale thermostat (Bussi et al. 2007) at 310 K (τ_*T*_ = 1.0 ps), followed by 5-step NPT equilibrations with increasing timesteps (*d*t=[1, 2, 5, 10, 20] fs) using the Berendsen barostat (Berendsen et al. 1984) at 1 bar (τ_*P*_ = 12.0 ps) with a compressibility of 3.0 10^−4^. During the initial two equilibration steps, position restraints were applied to the protein backbone (BB) beads. Additionally, the phosphate (PO4) beads of POPC lipids were weakly restrained along the z-axis to prevent out-of-plane fluctuations of the lipids. The position restraints on BB and PO4 beads were gradually removed during the last NPT equilibration step. Production runs were carried out using a 20 fs timestep with the velocity-rescale thermostat (Bussi et al. 2007) and the Parrinello-Rahman barostat (Parrinello and Rahman 1981).

To study the curvature sensing by IDR-containing proteins, buckled membrane systems were built by embedding the four different protein molecules in a pre-equilibrated POPC bilayer buckle. The initial shape of the membrane buckle was obtained using Lipid-Wrapper (Durrant and Amaro 2014) and equilibrated to form a continuous carpet-like folded membrane along the *x*-axis in a fixed *xy*-plane (57 ×28 nm^2^) under periodic boundary conditions. The proteins were initially placed in a region with low mean curvature, *i.e*., *H*(*x, y*) ≃ 0 nm^−1^ followed by system equilibration (NPT conditions: 1 bar; 310 K; anisotropic barostat) and production runs (up to 10 μs) to quantify curvature preferences.

To study protein-induced spontaneous budding in simulations, we used flat asymmetric bilayers varying the copy number (*n*_Prot_ = [1, 3, 6, 9] molecules) of different membrane proteins. Proteins were arranged in a square grid, ensuring each protein was separated from its nearest neighbor by approximately 10 nm. We varied relative asymmetries of POPC bilayers (∼2%, ∼7% and ∼12%) by changing the Δ*N* = *N*_upper_ *-N*_lower_, *i.e*., the difference in the number of lipids between the upper and the lower leaflets (Δ*N* = [0, 100, 300, 500, 700]) using `insane.py` (Wassenaar et al. 2015). Production runs were performed for three replicates (NPT conditions: 1 bar; 310 K; semi-isotropic barostat) for 5 μs.

### D. Rescaling protein-protein interactions

To reproduce the correct ensemble of IDR conformations sampled using the Martini models, the protein-protein LJ interactions were rescaled following a previously implemented method (Benayad et al. 2020). The parameter *α* scales the well-depth (ϵ) of the protein-protein LJ potential, ϵ_*α*_ = ϵ_0_ +*α* (ϵ_original_ ϵ_0_), such that *α* =0 corresponds to repulsion-dominated interaction in the Martini model, ϵ_0_ =2 kJ/mol, and *α* =1 recovers the full interaction of the Martini force field, ϵ_0_ = ϵ_original_. This procedure was previously used to model IDR dynamics, formation of phase-separated condensates (Benayad et al. 2020) and demonstrate the role of ubiquitination in RHD-clustering on bilayers (González et al. 2023). Multiple simulations using rescaled PPI at various *α*-values were performed to optimize it for the IDR of FAM134B. The conformational landscape of IDR ensembles in solution was then compared with atomistic MD simulations to obtain an optimal value (*α* = 0.6). To model the interactions of structured and intrinsically disordered segments within the same polypeptide chain, we used hybrid scaling with two different ϵ_*α*_ for self-interactions. ϵ_1_ modeled the LJ interactions within the folded/structured region (*e.g*., KALP_25_/RHD), and ϵ_0.6_ for interactions within the IDR. LJ parameters for cross-interactions between folded or structured and IDR segments were obtained using Lorentz-Berthelot combination rules, 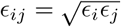 (Boda and Henderson 2008). The IDR interactions with all other molecules (lipids and solvent) were unchanged corresponding to ϵ_1.0_ (Thomasen et al. 2023).

### E. Backmapping coarse-grained structures

CG structures of IDR ensembles were back-mapped to all-atom resolution for computing ensemble properties of the IDR before comparisons with atomistic MD simulations. Atomistic models were obtained using `backward.py` script (Wassenaar et al. 2014) with appropriate CHARMM36m topology, and energy minimized using the default protocol. Atomistic models of KALP_25_– IDR and RHD–IDR from CGMD simulations were also obtained for computing hydrodynamic properties.

### F. IDR ensemble properties

The IDR from various MD simulations in their varied molecular and structural contexts were characterized by computing their ensemble properties. The radius of gyration (*R*_*g*_), end-to-end distances (*R*_*e*_), residue-wise local secondary structure assignments, and root mean square fluctuations (RMSF) for the MD trajectories were computed using the gmx functions (implemented in GRO-MACS package) `gyrate, distance, do_dssp`, and `rmsf`, respectively. The local secondary structure probability of the IDR was obtained by counting the relative frequency of *α*-helix (*P*_H_), β-strand (*P*_S_) or coil-like (*P*_C_) conformations assigned to each residue over the entire atomistic MD trajectory.

Free energy landscapes for back-mapped IDR ensembles in solution and membrane-anchored configurations were obtained from the sampled distributions *P*(*R*_*g*_, *R*_*e*_) as F (*R*_*g*_, *R*_*e*_) = ln *P*(*R*_*g*_, *R*_*e*_) in units of kcal mol^−1^. Free energy surfaces were computed on concatenated equilibrium MD trajectories of multiple replicates obtained using the `polystat` function of GROMACS.

The hydrodynamic properties of the IDR, such as asphericity, translational hydrodynamic radius (*R*_*T*_), and rotational hydrodynamic radius (*R*_*R*_), for the back-mapped MD trajectories of the IDR in solution, tethered to KALP_25_ and RHD were estimated with the HullRad (Fleming and Fleming 2018). Moreover, the convex hulls were visualized using `Display_hull3.py` script (Fleming and Fleming 2018).

### G. Contact maps

Contact maps for IDR-IDR, IDR–KALP_25_, and IDR–RHD interactions were obtained using *in-house* scripts implementing MDAnalysis (Michaud-Agrawal et al. 2011). The residue-wise interactions between two groups, A and B, were estimated for each frame of the trajectory such that AB_cnts_ =Σ _*i∈A j∈B σ*_( |*r*_*ij*_|), where the summation extends over heavy-atom positions of interacting residues (i, j), and *σ*(|*r*_*ij*_|) = 1 − [0.5 − 0.5(tanh((|*r*_*ij*_| − *r*_*c*_)/a))], is a smooth sigmoidal counting function that limits contacts below a cut-off distance, *r*_*c*_. We used *r*_*c*_ ≤ 5 Å with a = 0.5 Å for atomistic systems and *r*_*c*_ ≤ 10 Å with *a* = 1.0 Å for coarse-grained systems, respectively.

### H. Estimating membrane curvatures

Membrane shapes were characterized by quantifying local bilayer properties using a modified version of MemCurv (https://github.com/bio-phys/MemCurv) (Bhaskara et al. 2019). The MD trajectories were post-processed to align all frames such that the protein molecule was centered and oriented parallel to the *x* axis 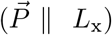. We used the long axis of the central amphipathic helix, (AH_L_) within the RHD to define its in-plane orientation 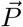. This ensured the removal of effects arising from the protein’s in-plane rotational and translational diffusion. The shape of the bilayer is characterized by a height function *h*(*x, y*) of the midplane using a Monge parametrization and approximated by an optimized 2D Fourier function as implemented in MemCurv (Bhaskara et al. 2019). The local height *h*(*x, y*) was then used to estimate the local shape operator **S**(*x, y*), which embodies all the curvature properties of the membrane surface. Local directional curvatures (*k*_1_ and *k*_2_), the mean curvature, *H* = Tr**(S)**/2, and the Gaussian curvature, *K*_G_ = det(**S**) are computed for the protein center-of-mass (COM) positions along the membrane by computing the eigenvalues, the trace and the determinant of the local shape operator **S**(*x, y*), as implemented in MemCurv, respectively. Computations were performed by fitting PO4 beads for every frame to obtain a height profile. The *H*(*x, y*) could then be computed for 40 ×40 grid (width = 1 nm) using MemCurv. *h*(*x, y*) and *h*(*x, y*) were averaged over 500 frames spanning the last 1 μs of the representative trajectories.

For buckled membranes, the lateral diffusion of the protein was tracked by measuring its COM positions along with local shape changes of the buckle, following the original version of MemCurv (Bhaskara et al. 2019). The local shape operator **S**(*x, y*) was calculated as previously described in order to extract values of *k*_1_, *k*_2_, *H*(*x, y*), and *K*_G_(*x, y*).

### I. Analysis of membrane budding and protein clustering

Several ordered parameters were used to monitor the spontaneous membrane budding of asymmetric bilayers containing different proteins. Sudden drastic changes in the time series of box width (*L*_*x*_) provided excellent readout on budding. The average waiting time, ⟨*t*⟩ = (*t*_1_ + *t*_2_ + *t*_3_)/3 for a 50% drop in *L*_*x*_ from 3 independent replicates were used to estimate the rate of budding *K* = 1/ ⟨*t*⟩ for each protein-membrane system. To ascertain the role of proteins and the associated shape transition into a bud-like geometry, the projection of the center-of-mass of the protein BB beads and the PO4 beads of both leaflets along the z axis was monitored. This provides a way to track the proteins along the bud shape relative to the highest and lowest points during the budding transition. Further, in MD simulations containing multiple protein copies, a pair-wise inter-protein distance matrix was defined using COM positions of individual proteins along the *xy* plane for each simulation frame. Hierarchical clustering of this matrix with single-linkage and cut-off distance of 10 nm was used to estimate the total number of clusters and the size of the largest protein cluster in each frame of the trajectory. Monitoring the time series of cluster sizes provided information on the role of IDR-mediated protein clustering and associated shape changes in membranes. Further, contact maps of residue-wise IDR-IDR interactions were also estimated for each inter-protein pair and averaged over the entire trajectory to map the most frequent IDR contact sites during budding.

## V. ACKNOWLEDGMENTS

This work is partially supported by the Cluster project EnABLE, funded by the Hessian Ministry for Science and Arts, and the CRC project on Selective Autophagy, DFG Project-ID 259130777-SFB1177 (to I.D., G.H., and R.M.B). S.A.P-C. and K.L. acknowledges funding from the EnABLE cluster and the SFB-1177 consortia, respectively. I.D. is also supported by the European Research Council (grant ER-REMODEL). G.H. thanks the Max Planck Society for support. We thank the Center for Supercomputing, Goethe University Frankfurt, for computing time on the Goethe-HLR cluster. We also thank Arghya Dutta and Lukas Stelzl for the fruitful discussions and David Krause for system administration.

## VI. CONFLICT OF INTEREST

The authors declare no competing interests.

## VII. AUTHOR CONTRIBUTIONS

R.M.B. conceived and designed the research based on discussions with G.H. S.A.P-C. developed the methodology, performed all modeling, simulations, and analyses with assistance from K.L. and support from R.M.B. I.D. and G.H. analyzed and interpreted the data, providing financial and technical support, respectively. S.A.P-C. and R.M.B. wrote the initial draft with input from all the authors.

## VIII. DATA AVAILABILITY STATEMENT

The data that support the findings of this study are available from the corresponding author upon reasonable request.

## X. SUPPLEMENTARY INFORMATION

Supplementary Figures (S1–S14), Tables (S1–S6), and Movie legends are provided in a separate PDF, along with Movie files SM1–SM6 (MP4).

## Notes

### Competing Interest Statement

The authors have declared no competing interest.

