## Supplementary Data File for "Intrinsically disordered region amplifies membrane remodeling to augment selective ER-phagy"

### Supplementary Information File:
### Intrinsically disordered region amplifies membrane remodeling to augment selective ER-phagy

Sergio Alejandro Poveda-Cuevas,[1,2] Kateryna Lohachova,[1,2] Ivan Dikic,[1,2] Gerhard Hummer,[3,4] and Ramachandra M. Bhaskara[1,2,a)]

[1)] *Goethe University Frankfurt, Institute of Biochemistry II, University Hospital Building 75, Theodor-Stern-Kai 7, 60590 Frankfurt (Main), Germany*

[2)] *Buchmann Institute for Molecular Life Sciences, Riedberg Campus, Max-von-Laue Str. 15, 60438 Frankfurt (Main), Germany*

[3)] *Max Planck Institute of Biophysics, Department of Theoretical Biophysics, Max-von-Laue Str. 3, 60438 Frankfurt (Main), Germany*

[4)] *Goethe University Frankfurt, Department of Physics, Institute of Biophysics, Max-von-Laue Str. 1, 60438 Frankfurt (Main), Germany*

(Dated: 28 March 2024)

This document contains the following sections:

1. **Supplementary Figures** (**S1-S14**)
2. **Supplementary Tables** (**S1-S6**)
3. **Supplementary Movie legends** (**SM1–SM6**)
3. **Supplementary References**

[a)] Electronic mail:

### I. SUPPLEMENTARY FIGURES

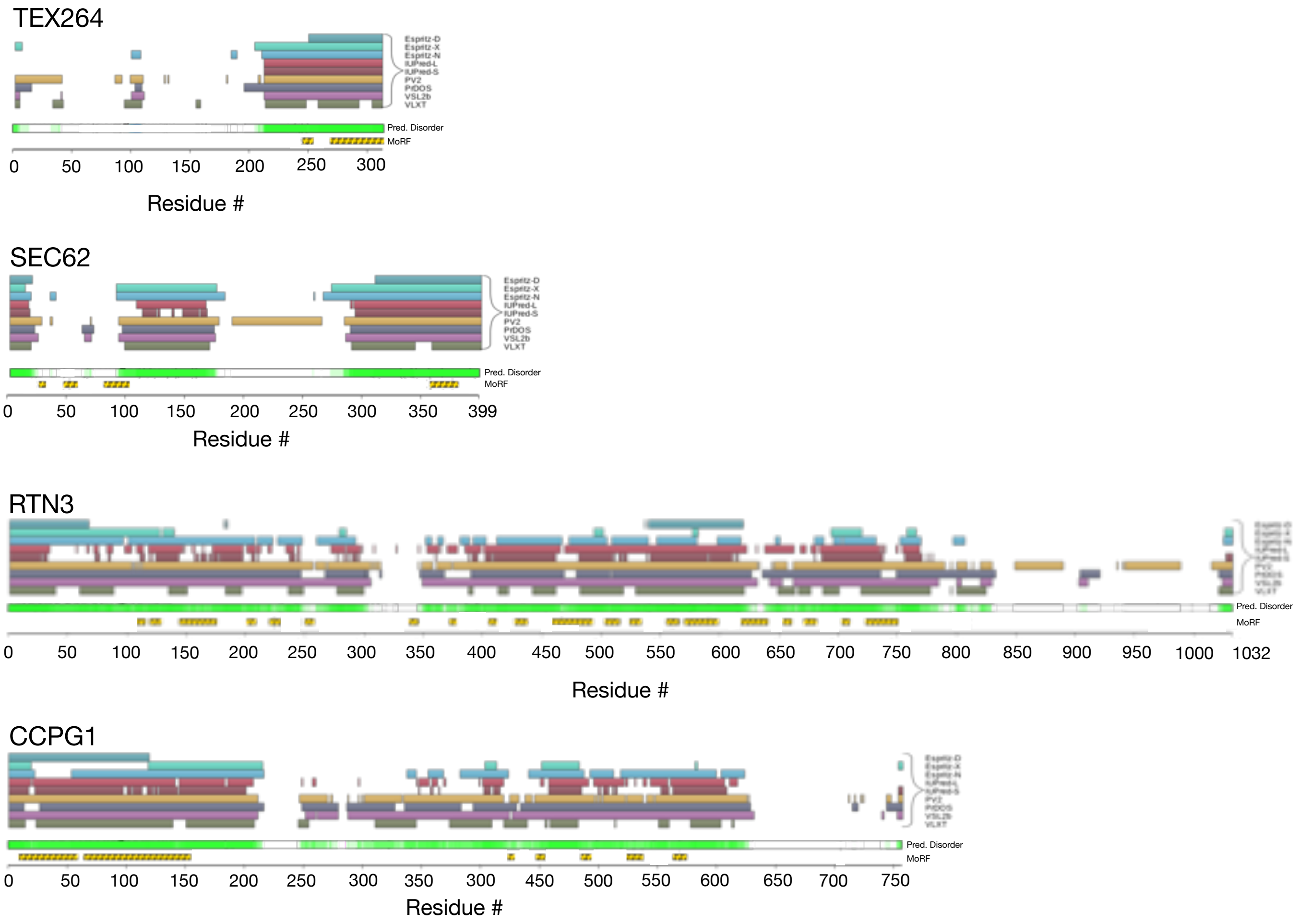


FIG. S1. **IDRs in human ER-phagy receptors.** IDRs of human TEX264, SEC62, RTN3, and CCPG1 were annotated using $D^2P^2$ (Oates et al. 2012), a consensus sequence-based IDR prediction method using nine different algorithms. N- and C-terminal IDR segments (green), along with predictions of molecular recognition feature (MoRF) based on ANCHOR predictions of protein-binding sites, are annotated. All known LIRs are housed within the cytosolic IDRs. TEX264 and SEC62 contain LIR sites in their C-terminal IDRs, whereas RTN3 and CCPG1 have LIRs in their N-terminal IDR segments.

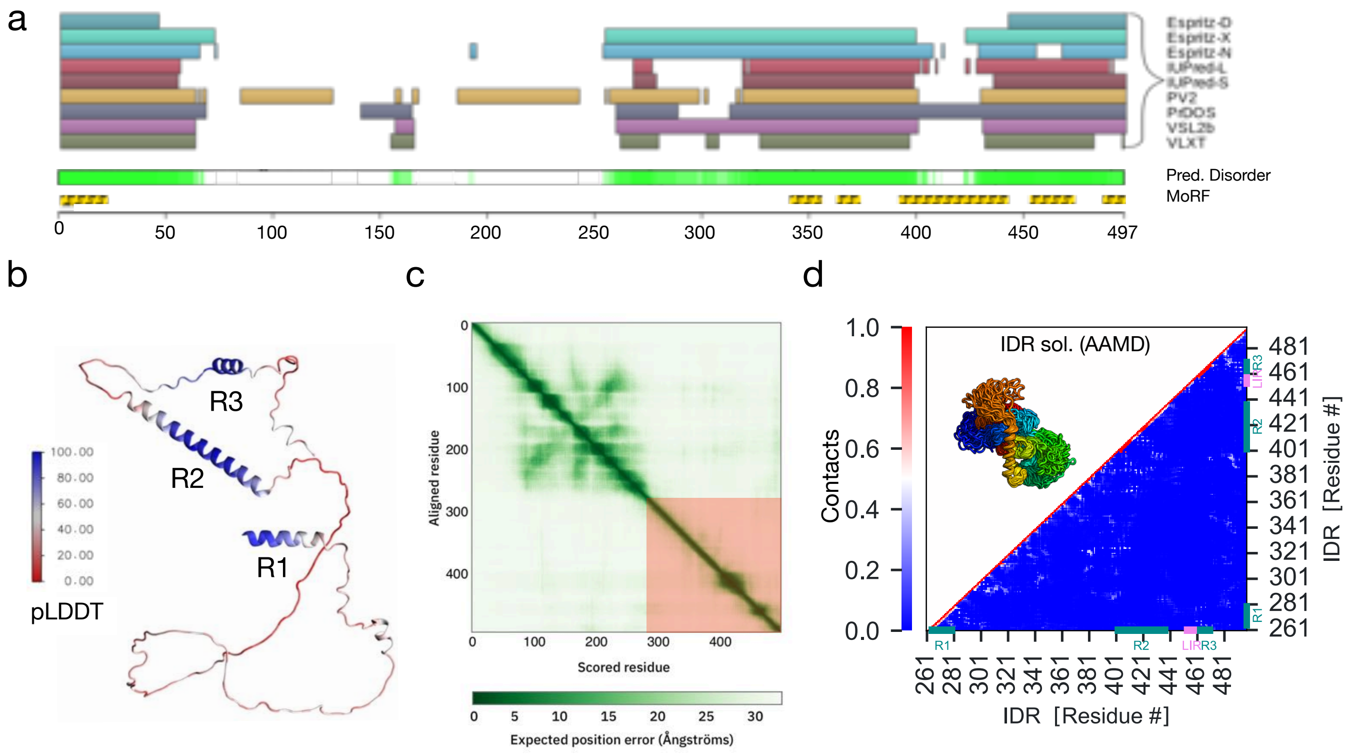


FIG. S2. **Sequence and structure of FAM134B–IDR.** (**a**) IDRs of FAM134B were predicted and annotated using D$^2$P$^2$ (Oates et al. 2012). (**b**) AF model (Jumper et al. 2021) of the C-terminal IDR (261–497) shows three helical regions with high confidence (R1, R2, and R3 with pLDDT≥ 70). (**c**) Predicted aligned error (PAE) for residue pairs $i, j$ corresponding to the C-terminal IDR (red box) and the RHD indicate uncertainty in their relative positions and orientations within the full-length AF model. (**d**) Residue-wise contact map of the FAM134B–IDR from atomistic MD simulations.

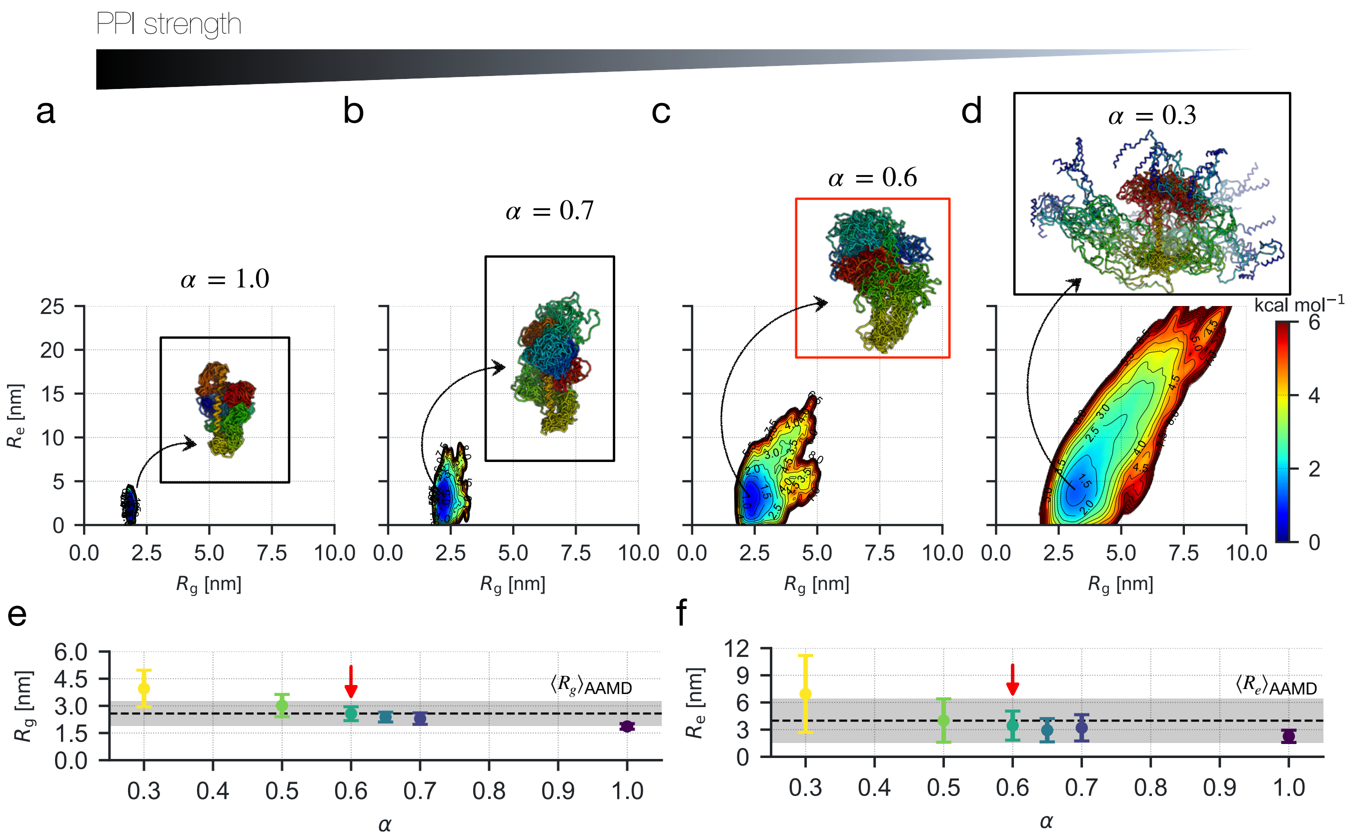


FIG. S3. **Scaling PPI strength is essential for effective coarse-graining of FAM134B-IDR.** (**a**–**d**) Free energy landscapes IDR ensembles in solution from the sampled distributions $p(R_g, R_e)$ using the scaled Martini models at $\alpha$ = [1.0, 0.7, 0.6, 0.3]. Insets display the ensemble of 30 representative structures of the IDR (rainbow colors) sampled in solution for each case. Average values of (**e**) $R_g$ and (**f**) $R_e$ (mean ± SD) of IDR-ensembles simulated by varying $\alpha$-scaling. The dashed line and the shaded region show corresponding mean values and standard deviations for structures sampled using CHARMM36m force-field.

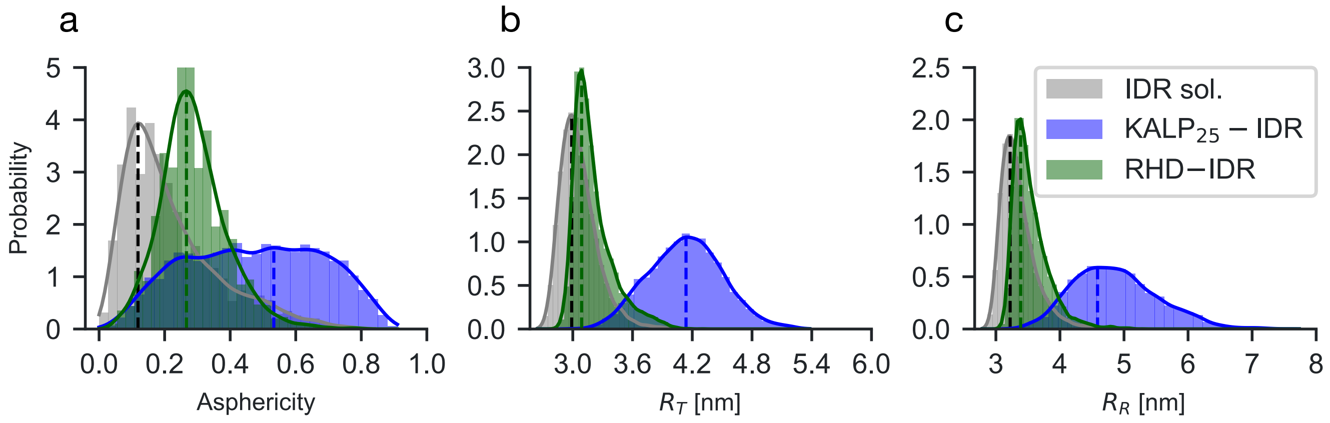


FIG. S4. **Hydrodynamic properties of FAM134B-IDR.** Distributions of (**a**) asphericity, (**b**) translational hydrodynamic radius ($R_T$) and (**c**) rotational hydrodynamic radius ($R_R$) of the FAM134B-IDR, in solution (grey), anchored to $KALP_{25}$ (blue), and anchored to RHD (green), respectively. Histograms and density estimates are computed for backmapped all-atom IDR structures sampled at 1-ns intervals over 5 $\mu$s coarse-grained MD trajectories using HullRad program (Fleming and Fleming 2018).

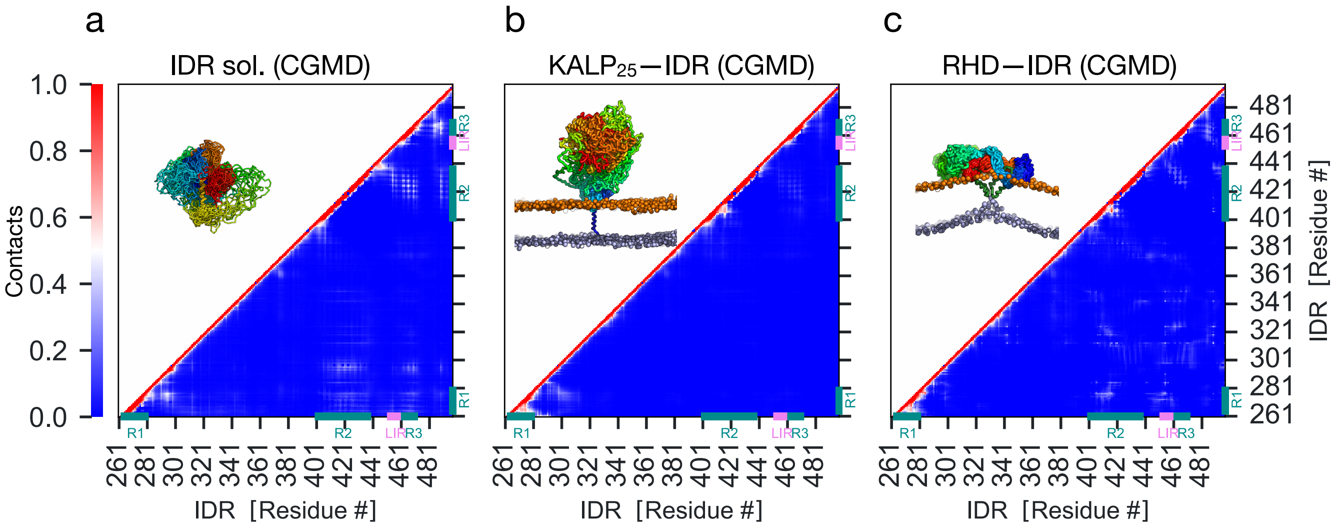


FIG. S5. **Context-dependent structure and interactions of FAM134B-IDR.** Intra-molecular pair-wise residue interactions averaged over the IDR ensemble sampled from coarse-grained MD simulations (5 $\mu$s runs; 3 replicates) (**a**) in dilute solution, (**b**) anchored to $KALP_{25}$, and (**c**) anchored to RHD, respectively. Regions of the IDR with residual helical structure (R1, R2, and R3) and the LIR motif are highlighted in cyan and purple, respectively. The upper triangle shows ensembles of 30 representative structures (rainbow-colored).

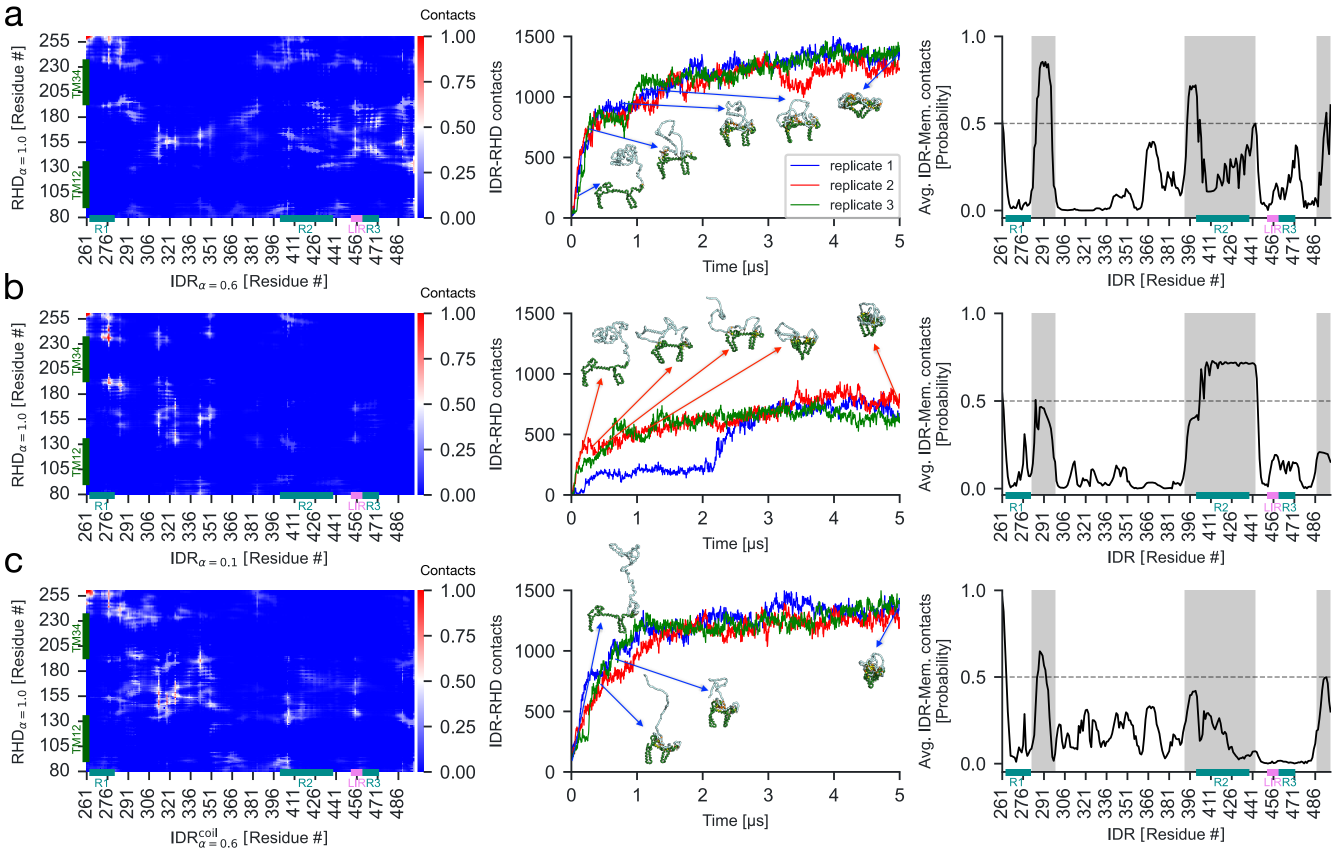


FIG. S6. **IDR interactions with the RHD and the membrane.** Three distinct RHD–IDR systems with altered scaling parameters: (**a**) Canonical optimized RHD–IDR ($\alpha = 0.6$), (**b**) RHD–IDR with reduced PPI ($\alpha = 0.1$), and (**c**) RHD–IDR with no residual helical structure, were modeled and simulated using adapted Martini model to study the influence of IDR-mediated interactions on the conformational ensemble. (**Left**) Pair-wise residue contact maps for IDR–RHD interactions averaged over $3 \times 5$ $\mu$s coarse-grained MD trajectories. Helical segments R1, R2, and R3, and the LIR motif of the IDR and the TM hairpins of the RHD are highlighted to show interactions. (**Middle**) Time series of IDR–RHD contacts characterize intermediate structures showing distinct stages of compaction, ultimately leading to RHD scaffolding in all three replicates. (**Right**) Average interactions of the IDR with the bilayer are quantified by contacts between BB and PO4 beads and mapped on IDR sequence. IDR segments with interaction probability $\geq 0.5$ (grey shaded) in the canonical RHD–IDR system correspond to residues flanking R2 and R1 stretches, respectively.

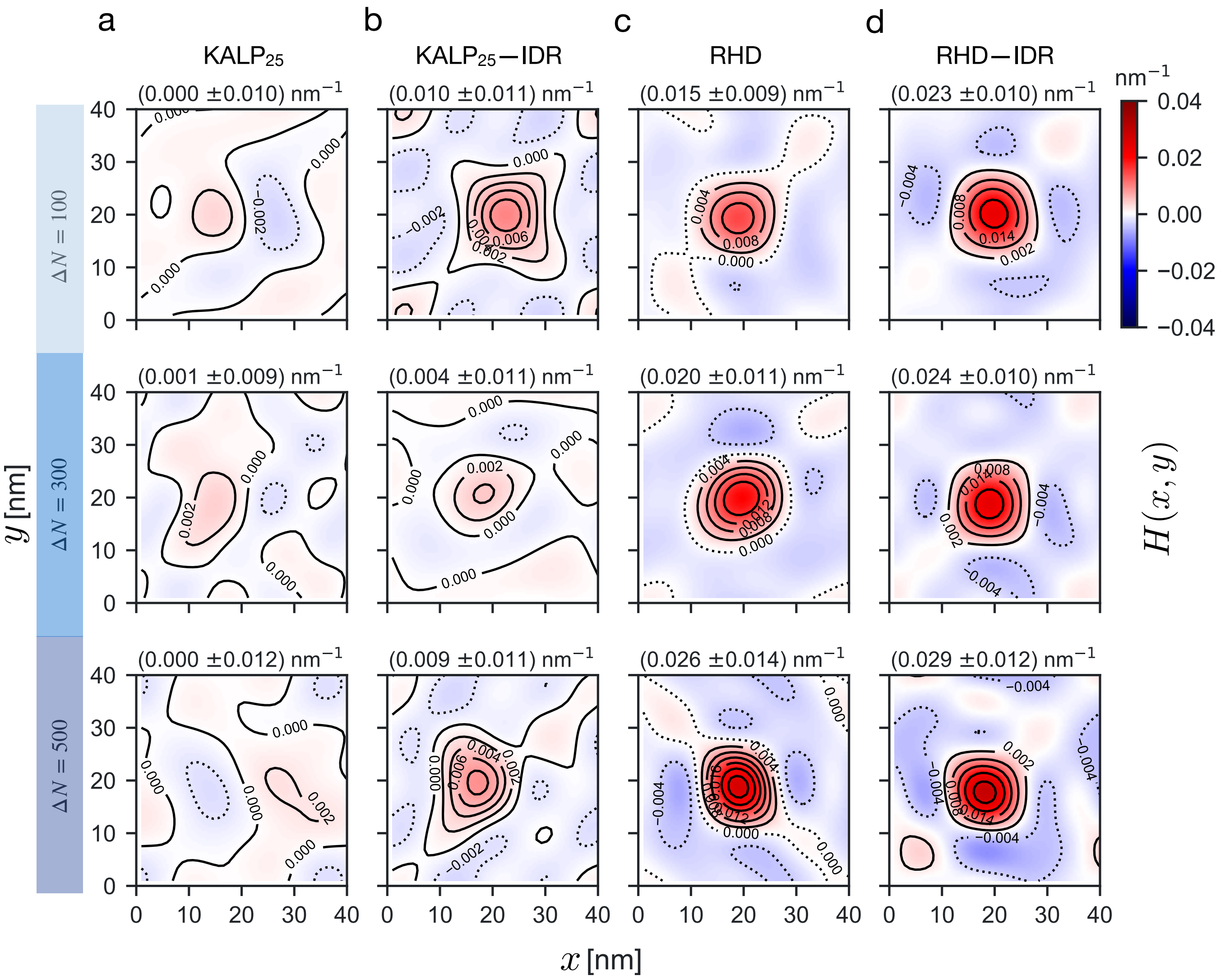


FIG. S7. **Effect of membrane asymmetry on protein-induced curvature.** Contour maps of local mean curvature field $H(x,y)$ induced by (**a**) $KALP_{25}$ (negative control), (**b**) $KALP_{25}$–IDR (**c**) RHD (positive control), and (**d**) RHD–IDR, respectively from flat asymmetric bilayers. Top-to-bottom panels show counter maps at increasing values of membrane asymmetry, $\Delta N = [100,\ 300,\ 500]$. The maximum value of curvature fields $\langle H_{\max} \rangle$ and $\pm$ SD is shown at the top. $H(x,y)$ are computed by fitting 500 individual frames at 2-ns intervals over the last 1 $\mu$s of representative trajectories.

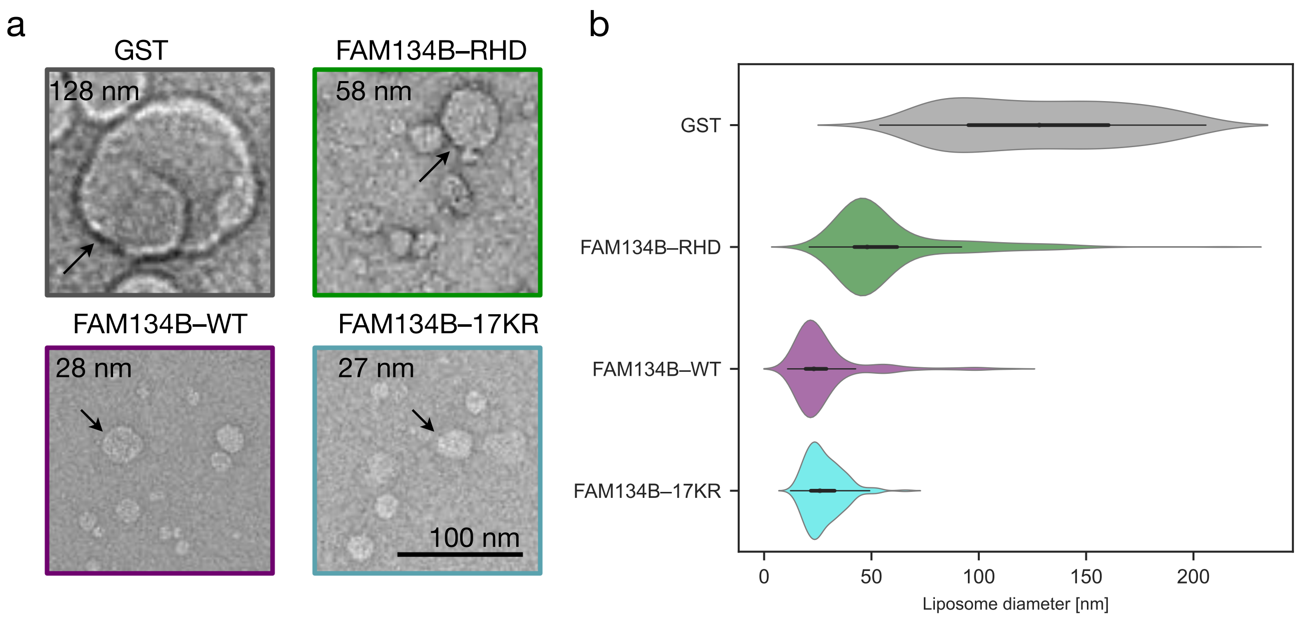


FIG. S8. **IDRs amplify *in vitro* liposome remodeling.** (**a**) Representative negative-stain transmission electron micrographs (re-scaled and cropped) comparing 200 nm liposomes remodeled by GST (negative control), FAM134B–RHD (RHD), full-length FAM134B–WT ($\simeq$ RHD–IDR), and FAM134B–17KR (18 K→R mutations in the RHD). (**b**) Violin plots of proteoliposome size distributions ($n = 300$ each) display the median value (black dot) and the interquartile range (black-shaded region), along with mirrored densities on either side (colored). Liposome remodeling data for FAM134B-RHD (without IDR) and RHDs with intact IDRs (WT, 17KR) are adapted from Fig. 6d, 6j in Bhaskara et al. (2019) and Extended Data Figs. 7d-e in González et al. (2023), respectively.

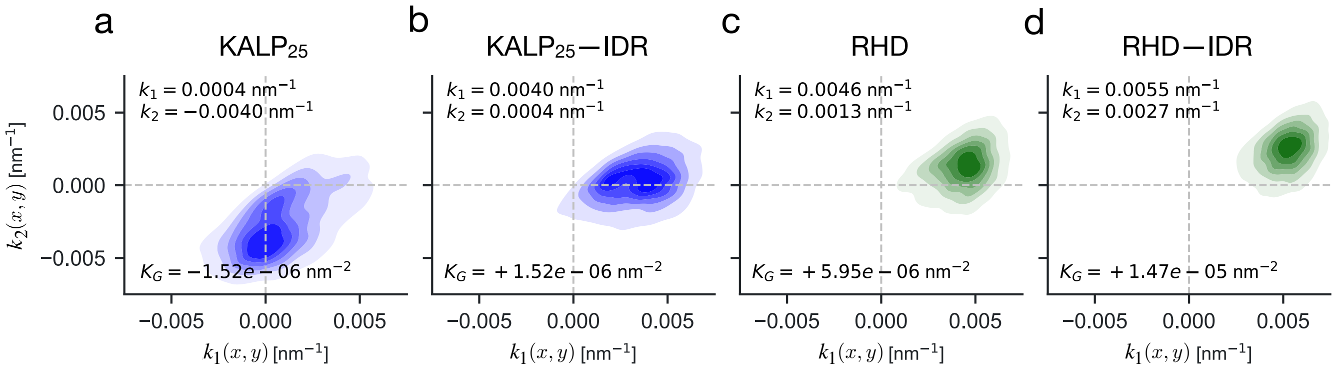


FIG. S9. **IDRs sense directional curvatures.** 2D-histograms (density indicated by color intensity) of the principal curvatures $k_1$ and $k_2$ sampled by (**a**) KALP$_{25}$ (negative control), (**b**) KALP$_{25}$–IDR (**c**) RHD (positive control), and (**d**) RHD–IDR embedded in membrane buckles (see Methods). KALP$_{25}$–IDR senses the top of the buckle, regions with tubular geometry ($k_1 > 0$, $k_2 \simeq 0$, and $K_G(x, y) \simeq 0$) as opposed to the free KALP$_{25}$-peptide, which prefers saddle-like structures ($k_1 \simeq 0$, $k_2 < 0$, and $K_G(x, y) \simeq 0$). IDRs anchored to RHDs perturb the buckle and prefer regions resembling ellipsoid vesicles with bi-directional curvatures ($k_1 > 0$, $k_2 > 1$, and $K_G(x, y) > 0$) at the top of the buckle.

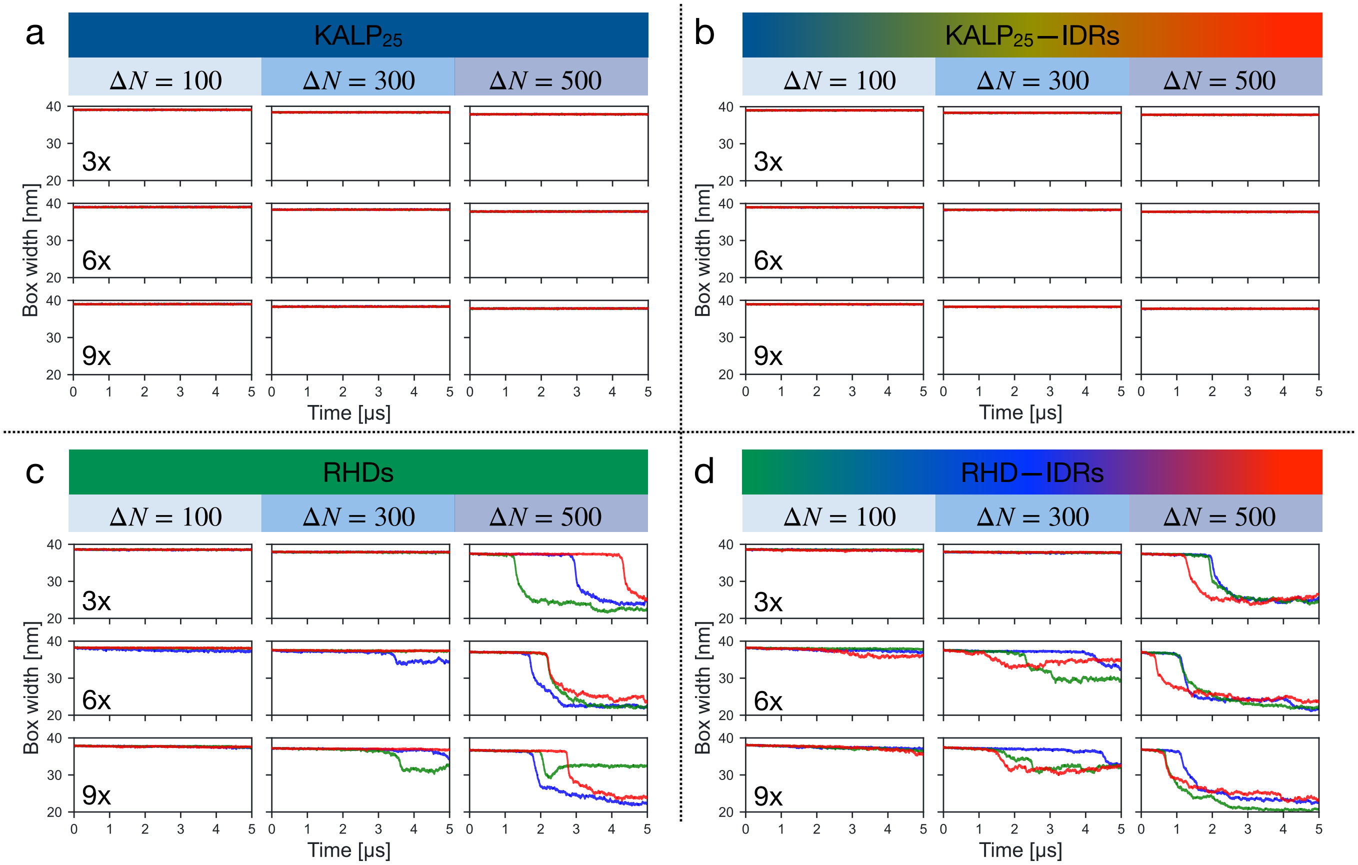


FIG. S10. **RHD–IDRs accelerate spontaneous membrane budding.** Time series of lateral box width $L_x$ report on spontaneous budding events in coarse-grained MD simulations initiated from asymmetric bilayers (see Methods). Systems containing different copies ($n_{\mathrm{Prot}}$ = [3, 6, 9]; top-to-bottom) of (**a**) $\mathrm{KALP}_{25}$ (negative control), (**b**) $\mathrm{KALP}_{25}$–IDR (**c**) RHD (positive control), and (**d**) RHD–IDR. Protein molecules were embedded in flat asymmetric bilayers ($\Delta N$ = [100, 300, 500]; left-to-right) to study spontaneous budding events. MD trajectories for each system were simulated (3 × 5 $\mu$s runs; red, blue, and green curves) to measure the kinetics of budding. Waiting times for budding were estimated as the time taken for a 50% drop in the box width, $L_x$.

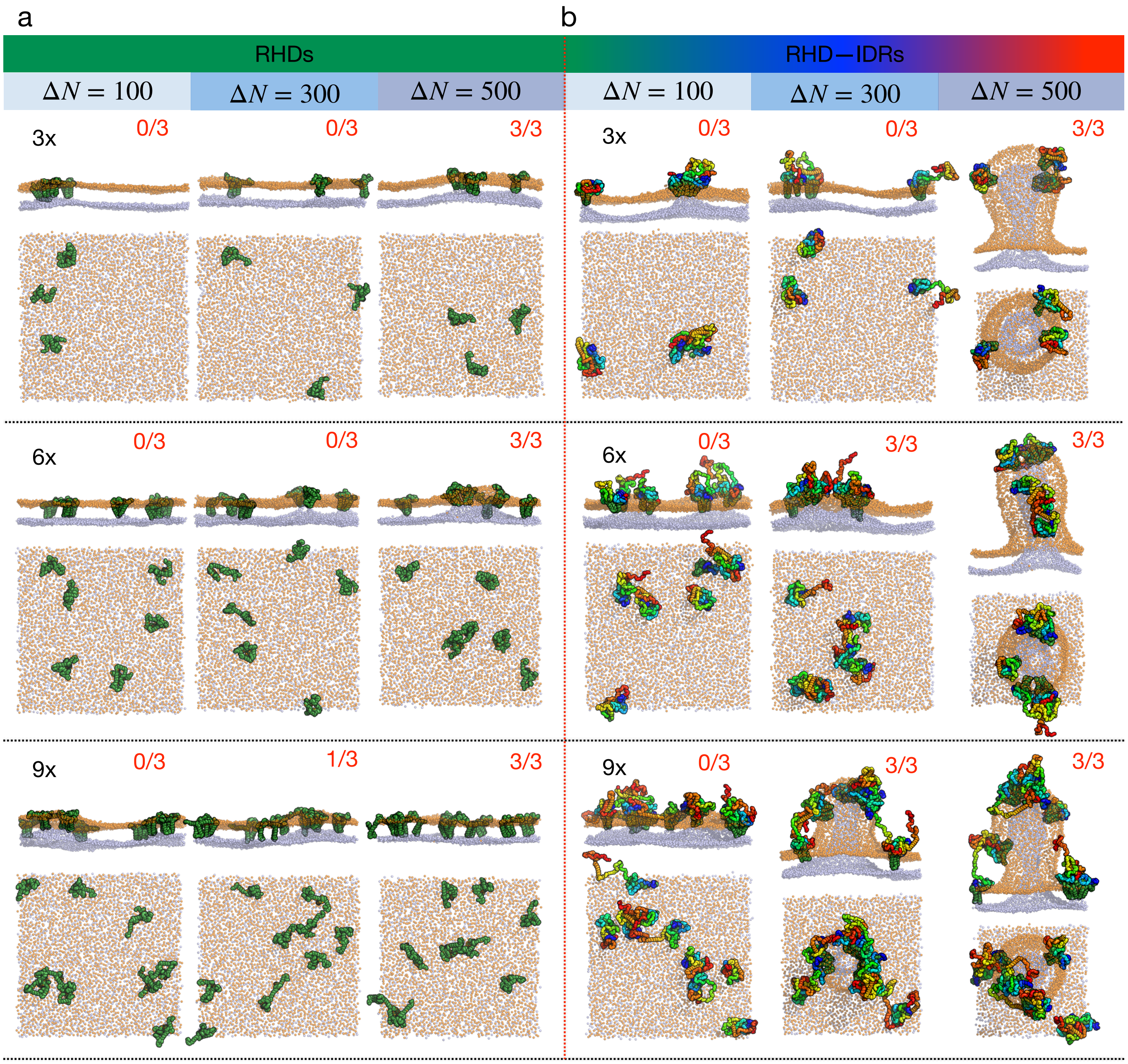


FIG. S11. **IDRs intensify RHD-mediated membrane budding.** Representative snapshots showing the side- and top-views of protein-membrane systems at the end of 2 $\mu$s MD simulations. They show asymmetric bilayers ($\Delta N$ = [100, 300, 500]) embedded with multiple copies ($n_{\mathrm{Prot}}$ = [3, 6, 9]) of (**a**) RHD or (**b**) RHD–IDR molecules. PO4 beads (orange spheres) display bilayer shape changes induced by RHDs (green) and IDRs (rainbow). The number of spontaneous budding events observed on a longer time-scale (5 $\mu$s) in 3 independent replicates are denoted (red fraction).

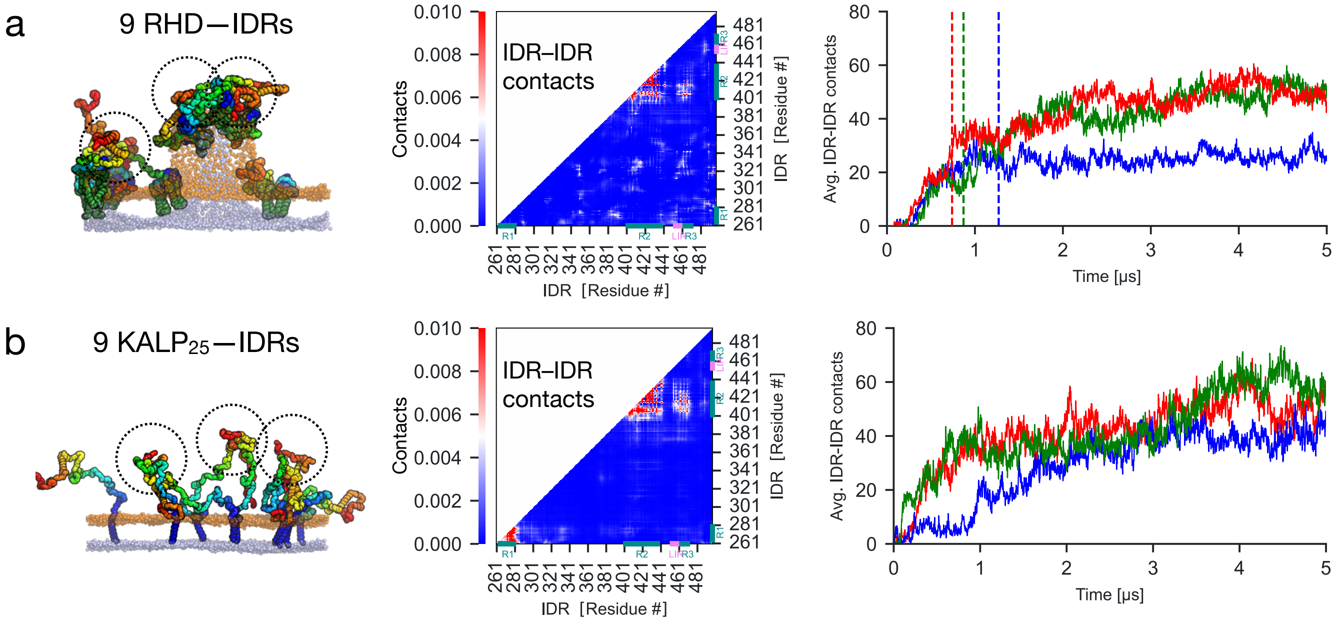


FIG. S12. **IDR interactions mediate membrane protein clustering.** Inter-molecular IDR–IDR contacts monitored in MD simulations containing 9 (**a**) RHD–IDR, and (**b**) KALP$_{25}$–IDR molecules influence protein clustering. (**Left**) IDRs anchored to RHDs mediate solution-phase interactions driving protein clustering, which, in turn, nucleate membrane buds spontaneously. (**Middle**) Residue-wise decomposition of IDR-IDR contacts shows interaction hotspots (red regions) corresponding to R2 and R3. (**Right**) A sudden increase in inter-molecular IDR-IDR contacts of RHD–IDRs $\geq$ 20 coincides with bud formation (dashed lines) in all three replicates (red, blue, green curves). Although KALP$_{25}$–IDRs display IDR-mediated inter-molecular contacts and result in clustering, they are insufficient to nucleate buds on the MD timescale.

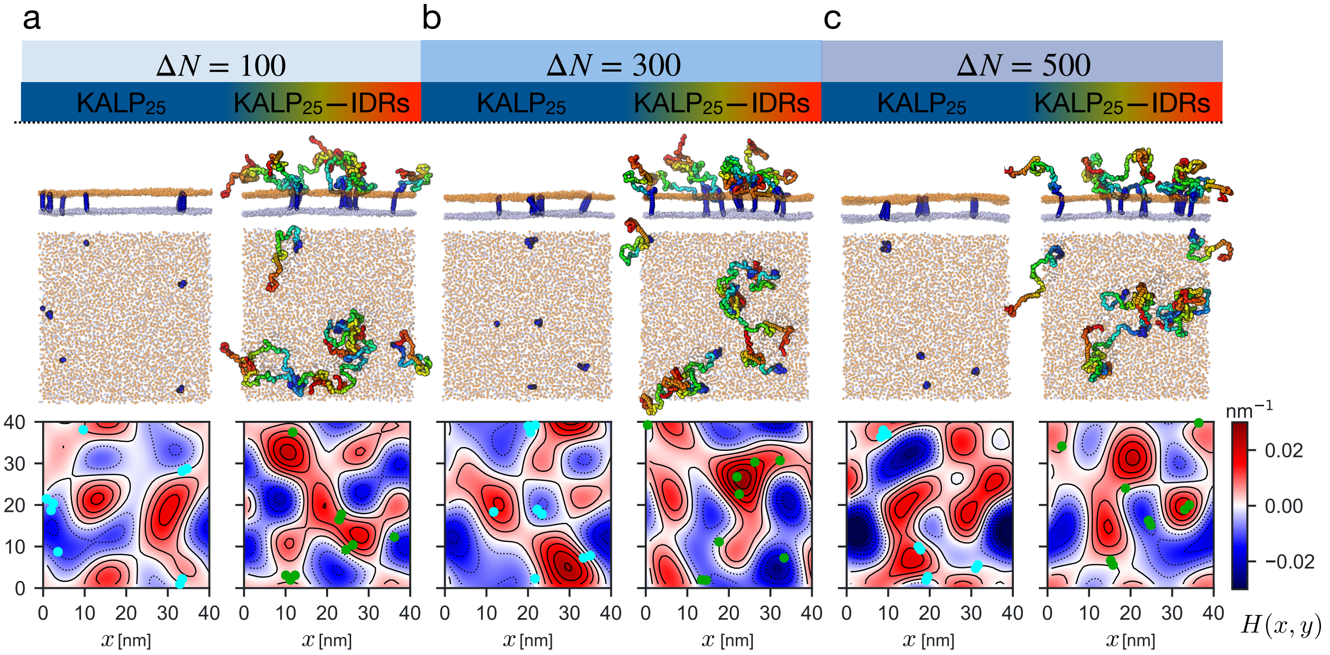


FIG. S13. **IDR-mediated contacts intensify local membrane curvature induction.** Comparison of MD simulations containing 9 KALP$_{25}$ and KALP$_{25}$–IDR molecules with (**a–c**) increasing membrane asymmetries. (**Top**) Snapshots at the end 5 $\mu$s show the prevalence of inter-molecular interactions mediated by the IDRs. (**Bottom**) Curvature contour maps of asymmetric bilayers at 5 $\mu$s. KALP$_{25}$ peptides oligomerize (cyan dots) but are inefficient in altering local membrane curvature protein even at high asymmetries. KALP$_{25}$–IDRs driven by their inter-molecular IDR contacts enhance the lifetime of the clusters and induce local bulging (green dots). They induce strong membrane bulges, which grow in intensity with increasing membrane asymmetry (dark red contour lines).

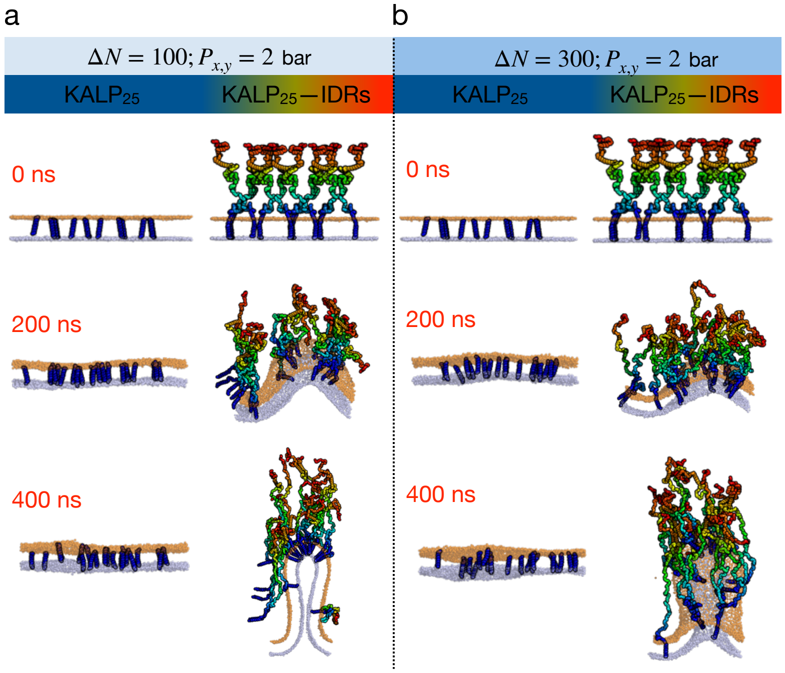


FIG. S14. **IDRs deform flat bilayers upon crowding.** (**a**–**b**) Asymmetric bilayers containing 20 $KALP_{25}$ peptides or KALP–IDR molecules were subjected to slight lateral compression $P_{xy} = 2$ bar. IDR-IDR interactions and crowding on one side of the membrane result in lateral bilayer stress, deforming the metastable flat asymmetric bilayers into buds. Asymmetric bilayers containing only $KALP_{25}$ peptides resist buckling and remain flat.

#### II. SUPPLEMENTARY TABLES

TABLE S1. **IDRs of human ER-phagy receptors.** Summary of IDR predictions using $D^2P^2$ (Oates et al. 2012) on well-known human ER-phagy receptors (UniProt accession codes). Cytosolic N-terminal or C-terminal IDRs containing the LIR motif (*) are mapped and used to annotate IDRs of their homologous proteins.

| Receptor | UnitProt accession code | Protein size | N-ter IDR | C-ter IDR | IDR length |
|---|---|---|---|---|---|
| FAM134B | Q9H6L5 | 497 | 1 – 92 | 261 – 497* | 237 |
| TEX264 | Q9Y6I9 | 313 | – | 213 – 313* | 100 |
| SEC62 | Q99442 | 399 | 108 – 167 | 290 – 399* | 110 |
| RTN3 | O95197 | 1032 | 1 – 768* | – | 768 |
| CCPG1 | Q9ULG6 | 757 | 1 – 208* | 250 – 599 | 208 |

TABLE S2. **MD simulations of FAM134B-IDR in solution.** Summary of atomistic and coarse-grained molecular systems used to model and simulate FAM134B-IDR in dilute solution (150 mM NaCl).

| Force Field | $\alpha$ | PBC | $\mathbf{v1}_{x,y,z}$ | $\mathbf{v2}_{y,z,x}$ | $\mathbf{v3}_{z,x,y}$ | Edge distance [nm] | $n_{\text{Prot}}$ | $N_{\text{Total}}$ | Water | $Na^+$ | $Cl^-$ | Time [$\mu$s] | Rep. |
|---|---|---|---|---|---|---|---|---|---|---|---|---|---|
| CHARMM36m (AA) | – | Dodecahedron | $\begin{bmatrix}2.3\\2.3\\1.6\end{bmatrix}$ | $\begin{bmatrix}0.0\\0.0\\0.0\end{bmatrix}$ | $\begin{bmatrix}0.0\\1.1\\1.1\end{bmatrix}$ | 4 | 1 | 807874 | 802839 | 770 | 737 | 1 | 2 |
| Martini 2.2 (CG) | 1.0 | Hexagonal | $\begin{bmatrix}3.5\\3.5\\2.5\end{bmatrix}$ | $\begin{bmatrix}0.0\\0.0\\0.0\end{bmatrix}$ | $\begin{bmatrix}0.0\\1.8\\1.8\end{bmatrix}$ | 2 | 1 | 256427 | 250421 | 2774 | 2741 | 10 | 2 |
| | 0.7 | Hexagonal | $\begin{bmatrix}3.5\\3.5\\2.5\end{bmatrix}$ | $\begin{bmatrix}0.0\\0.0\\0.0\end{bmatrix}$ | $\begin{bmatrix}0.0\\1.8\\1.8\end{bmatrix}$ | 2 | 1 | 256427 | 250421 | 2774 | 2741 | 10 | 2 |
| | 0.6 | Hexagonal | $\begin{bmatrix}3.5\\3.5\\2.5\end{bmatrix}$ | $\begin{bmatrix}0.0\\0.0\\0.0\end{bmatrix}$ | $\begin{bmatrix}0.0\\1.8\\1.8\end{bmatrix}$ | 2 | 1 | 256427 | 250421 | 2774 | 2741 | 10 | 2 |
| | 0.3 | Hexagonal | $\begin{bmatrix}3.5\\3.5\\2.5\end{bmatrix}$ | $\begin{bmatrix}0.0\\0.0\\0.0\end{bmatrix}$ | $\begin{bmatrix}0.0\\1.8\\1.8\end{bmatrix}$ | 2 | 1 | 256427 | 250421 | 2774 | 2741 | 10 | 2 |

TABLE S3. **Coarse-grained MD simulations of proteins embedded in flat bilayers.** Summary of coarse-grained MD simulations of various protein-membrane systems. Initial box dimensions ($L_x \times L_y \times L_z$), system size $N_{\text{Total}}$, number of protein molecules $n_{\text{Prot}}$, system size, number of POPC lipids in individual leaflets ($N_{\text{upper}}$, $N_{\text{lower}}$, and $\Delta N$) leaflets, number of water beads, ions, total simulation time, and the number of replicates for each system are provided.

| System | System size [$nm^3$] | $n_{\text{Prot}}$ | $N_{\text{Total}}$ | $\Delta N$ ($N_{\text{upper}}/N_{\text{lower}}$) | Water | $Na^+$ | $CL^-$ | Time [$\mu$s] | Rep. |
|---|---|---|---|---|---|---|---|---|---|
| $KALP_{25}$ + Mem. | 38 × 38 × 20 | 1 | 243345 | 0 (2398/2398) | 175732 | 2609 | 2613 | 5 | 3 |
| $KALP_{25}$–IDR + Mem. | 38 × 38 × 38 | 1 | 483215 | 0 (2397/2397) | 409895 | 5246 | 5218 | 5 | 3 |
| RHD + Mem. | 38 × 38 × 20 | 1 | 243037 | −14 (2384/2370) | 175588 | 2609 | 2617 | 5 | 3 |
| RHD–IDR + Mem. | 38 × 38 × 38 | 1 | 483219 | −14 (2389/2375) | 409915 | 5242 | 5218 | 5 | 3 |

TABLE S4. **Coarse-grained MD simulations in buckled membranes.** Summary of protein-membrane systems used for studying the curvature sensing properties of FAM134B-IDR. For each system, the membrane buckle is simulated in a periodic ($L_x \times L_y \times L_z$) with fixed x-y plane (see Methods). The number of protein molecules $n_{\text{Prot}}$, system size $N_{\text{Total}}$, total number of POPC molecules $N_{\text{lipids}}$, the number of water beads and ions, total simulation time, and the number of replicates are provided.

| System | System size [$nm^3$] | $n_{\text{Prot}}$ | $N_{\text{Total}}$ | $N_{\text{lipids}}$ | Water | $Na^+$ | $CL^-$ | Time [$\mu$s] | Rep. |
|---|---|---|---|---|---|---|---|---|---|
| Empty Mem. | 57 × 28 × 25 | – | 352868 | 5323 | 276343 | 3663 | 3663 | 10 | 1 |
| $KALP_{25}$+ Mem. | 57 × 28 × 25 | 1 | 351961 | 5296 | 275628 | 3719 | 3723 | 10 | 1 |
| $KALP_{25}$–IDR + Mem. | 57 × 28 × 25 | 1 | 416712 | 5295 | 337227 | 5072 | 5044 | 10 | 1 |
| RHD + Mem. | 57 × 28 × 25 | 1 | 345543 | 5212 | 269920 | 3719 | 3727 | 10 | 1 |
| RHD–IDR + Mem. | 57 × 28 × 25 | 1 | 413506 | 5208 | 334778 | 5068 | 5044 | 10 | 1 |

TABLE S5. **Coarse-grained MD simulations of asymmetric bilayers.** The table summarizes the various protein constructs used to study the effect of IDRs on membrane budding. We varied the number of proteins ($n$) and membrane asymmetry ($\Delta N$) to modulate the barrier for bud formation (see Methods) from flat metastable bilayers. Each system is represented by the periodic box, number of proteins $n_{\mathrm{Prot}}$, system size $N_{\mathrm{Total}}$, the bilayer asymmetry $\Delta N$, the number of lipids in the upper ($N_{\mathrm{upper}}$) and lower ($N_{\mathrm{lower}}$) leaflets, number of water/ions beads, total simulation time, and the number of replicates.

| System | System size [nm$^3$] | $n_{\mathrm{Prot}}$ | $N_{\mathrm{Total}}$ | $\Delta N$ ($N_{\mathrm{upper}}/N_{\mathrm{lower}}$) | Water | Na$^+$ | CL$^-$ | Time [$\mu$s] | Rep. |
|---|---|---|---|---|---|---|---|---|---|
| KALP$_{25}$ + Mem. | 38 × 38 × 20 | 1 | 242302 | 100 (2398/2298) | 175989 | 2609 | 2613 | 5 | 3 |
| | | 3 | 242253 | 100 (2392/2292) | 176002 | 2609 | 2621 | 5 | 3 |
| | | 6 | 242122 | 100 (2381/2281) | 176016 | 2609 | 2633 | 5 | 3 |
| | | 9 | 242119 | 100 (2373/2273) | 176080 | 2609 | 2645 | 5 | 3 |
| | | 20 | 241898 | 100 (2338/2238) | 176252 | 2609 | 2689 | 5 | 1 |
| | 38 × 38 × 20 | 1 | 240218 | 300 (2398/2098) | 176505 | 2609 | 2613 | 5 | 3 |
| | | 3 | 240171 | 300 (2392/2092) | 176520 | 2609 | 2621 | 5 | 3 |
| | | 6 | 240104 | 300 (2381/2081) | 176598 | 2609 | 2633 | 5 | 3 |
| | | 9 | 240053 | 300 (2373/2073) | 176614 | 2609 | 2645 | 5 | 3 |
| | | 20 | 239851 | 300 (2338/2038) | 176805 | 2609 | 2689 | 5 | 1 |
| | 38 × 38 × 20 | 1 | 238266 | 500 (2398/1898) | 177153 | 2609 | 2613 | 5 | 3 |
| | | 3 | 238205 | 500 (2392/1892) | 177154 | 2609 | 2621 | 5 | 3 |
| | | 6 | 238111 | 500 (2381/1881) | 177205 | 2609 | 2633 | 5 | 3 |
| | | 9 | 238117 | 500 (2373/1873) | 177278 | 2609 | 2645 | 5 | 3 |
| | | 20 | 237948 | 500 (2338/1838) | 177502 | 2609 | 2689 | 5 | 1 |
| KALP$_{25}$–IDR + Mem. | 38 × 38 × 38 | 1 | 460344 | 100 (2397/2297) | 388846 | 4985 | 4957 | 5 | 3 |
| | | 3 | 460713 | 100 (2392/2292) | 388221 | 5041 | 4957 | 5 | 3 |
| | | 6 | 461105 | 100 (2381/2281) | 387213 | 5125 | 4957 | 5 | 3 |
| | | 9 | 461718 | 100 (2377/2277) | 386244 | 5209 | 4957 | 5 | 3 |
| | | 20 | 463426 | 100 (2351/2251) | 382446 | 5517 | 4957 | 5 | 1 |
| | 38 × 38 × 38 | 1 | 458220 | 300 (2397/2097) | 389322 | 4985 | 4957 | 5 | 3 |
| | | 3 | 458571 | 300 (2392/2092) | 388679 | 5041 | 4957 | 5 | 3 |
| | | 6 | 458985 | 300 (2381/2081) | 387693 | 5125 | 4957 | 5 | 3 |
| | | 9 | 459598 | 300 (2377/2077) | 386724 | 5209 | 4957 | 5 | 3 |
| | | 20 | 461376 | 300 (2351/2051) | 382996 | 5517 | 4957 | 5 | 1 |
| | 38 × 38 × 38 | 1 | 456246 | 500 (2397/1897) | 389948 | 4985 | 4957 | 5 | 3 |
| | | 3 | 456602 | 500 (2392/1892) | 389310 | 5041 | 4957 | 5 | 3 |
| | | 6 | 457044 | 500 (2381/1881) | 388352 | 5125 | 4957 | 5 | 3 |
| | | 9 | 457592 | 500 (2377/1877) | 387318 | 5209 | 4957 | 5 | 3 |
| | | 20 | 459400 | 500 (2351/1851) | 383620 | 5517 | 4957 | 5 | 1 |
| RHD + Mem. | 38 × 38 × 20 | 1 | 241818 | 100 (2370/2270) | 175851 | 2609 | 2617 | 5 | 3 |
| | | 3 | 241649 | 100 (2321/2221) | 176098 | 2609 | 2633 | 5 | 3 |
| | | 6 | 241167 | 100 (2240/2140) | 177136 | 2609 | 2657 | 5 | 3 |
| | | 9 | 240373 | 100 (2159/2059) | 176460 | 2609 | 2681 | 5 | 3 |
| | 38 × 38 × 20 | 1 | 239853 | 300 (2370/2070) | 176486 | 2609 | 2617 | 5 | 3 |
| | | 3 | 239599 | 300 (2321/2021) | 176648 | 2609 | 2633 | 5 | 3 |
| | | 6 | 239268 | 300 (2240/1940) | 177136 | 2609 | 2657 | 5 | 3 |
| | | 9 | 238539 | 300 (2159/1859) | 177226 | 2609 | 2681 | 5 | 3 |
| | 38 × 38 × 20 | 1 | 237875 | 500 (2370/1870) | 177108 | 2609 | 2617 | 5 | 3 |
| | | 3 | 237705 | 500 (2321/1821) | 177354 | 2609 | 2633 | 5 | 3 |
| | | 6 | 237363 | 500 (2240/1740) | 177831 | 2609 | 2657 | 5 | 3 |
| | | 9 | 236765 | 500 (2159/1659) | 178052 | 2609 | 2681 | 5 | 3 |
| RHD–IDR + Mem. | 38 × 38 × 38 | 1 | 459097 | 100 (2373/2273) | 387849 | 4981 | 4957 | 5 | 3 |
| | | 3 | 459093 | 100 (2322/2222) | 387299 | 5029 | 4957 | 5 | 3 |
| | | 6 | 459328 | 100 (2239/2139) | 386884 | 5101 | 4957 | 5 | 3 |
| | | 9 | 459365 | 100 (2187/2087) | 385768 | 5173 | 4957 | 5 | 3 |
| | 38 × 38 × 38 | 1 | 457069 | 300 (2373/2073) | 388421 | 4981 | 4957 | 5 | 3 |
| | | 3 | 457172 | 300 (2322/2022) | 387978 | 5029 | 4957 | 5 | 3 |
| | | 6 | 457350 | 300 (2239/1939) | 387506 | 5101 | 4957 | 5 | 3 |
| | | 9 | 457471 | 300 (2187/1887) | 386429 | 5173 | 4957 | 5 | 3 |
| | 38 × 38 × 38 | 1 | 455204 | 500 (2373/1873) | 389156 | 4981 | 4957 | 5 | 3 |
| | | 3 | 455251 | 500 (2322/1802) | 388657 | 5029 | 4957 | 5 | 3 |
| | | 6 | 455509 | 500 (2239/1739) | 388265 | 5101 | 4957 | 5 | 3 |
| | | 9 | 455708 | 500 (2187/1687) | 387271 | 5173 | 4957 | 5 | 3 |

TABLE S6. **Kinetics of membrane budding from coarse-grained MD simulations.** Spontaneous budding events were monitored for each system (see Supplementary Table S5) by measuring waiting times ($t$) for 50% drop in box vector ($L_x$) from the initial value. The number of budding events for each system is represented as a fraction ($N_{\mathrm{buds}}/3$). The rate of budding $k = 1/\langle t \rangle$, where $\langle t \rangle = (t_1 + t_2 + t_3)/3$ and the acceleration factor $a = k_{\mathrm{system}}/k_{\mathrm{RHD}}$ are shown.

| System | $n_{\mathrm{Prot}}$ | $\triangle N$ | Asymm. [%] | $N_{\mathrm{Buds}}/3$ | $\langle t \rangle \pm$ SD [$\mu s$] | $k$ [$\mu s^{-1}$] | $a$ |
|---|---|---|---|---|---|---|---|
| RHD + Mem. | 1 | 100 | 2.15 | 0/3 | – | – | – |
| | 3 | | 2.20 | 0/3 | – | – | – |
| | 6 | | 2.28 | 0/3 | – | – | – |
| | 9 | | 2.37 | 0/3 | – | – | – |
| | 1 | 300 | 6.76 | 0/3 | – | – | – |
| | 3 | | 6.91 | 0/3 | – | – | – |
| | 6 | | 7.18 | 0/3 | – | – | – |
| | 9 | | 7.47 | 1/3 | 5 | 0.2 | 1 |
| | 1 | 500 | 11.79 | 0/3 | – | – | – |
| | 3 | | 12.07 | 3/3 | $2 \pm 2$ | 0.53 | 1 |
| | 6 | | 12.56 | 3/3 | $1.53 \pm 0.02$ | 0.65 | 1 |
| | 9 | | 13.10 | 3/3 | $1.3 \pm 0.5$ | 0.74 | 1 |
| RHD–IDR + Mem. | 1 | 100 | 2.15 | 0/3 | – | – | – |
| | 3 | | 2.20 | 0/3 | – | – | – |
| | 6 | | 2.28 | 0/3 | – | – | – |
| | 9 | | 2.37 | 0/3 | – | – | – |
| | 1 | 300 | 6.75 | 0/3 | – | – | – |
| | 3 | | 6.91 | 0/3 | – | – | – |
| | 6 | | 7.18 | 3/3 | $2 \pm 1$ | 0.57 | – |
| | 9 | | 7.45 | 3/3 | $2 \pm 2$ | 0.67 | 3.35 |
| | 1 | 500 | 11.77 | 0/3 | – | – | – |
| | 3 | | 12.06 | 3/3 | $1.8 \pm 0.6$ | 0.85 | 1.60 |
| | 6 | | 12.57 | 3/3 | $1.2 \pm 0.4$ | 0.87 | 1.65 |
| | 9 | | 13.07 | 3/3 | $0.7 \pm 0.3$ | 1.48 | 2.00 |

#### III. SUPPLEMENTARY MOVIE LEGENDS

- **Movie SM1: Atomistic MD simulations of FAM134B-IDR in solution.**
  The movie shows a 1 $\mu$s trajectory of FAM134B-IDR adopting a compact structure in a dilute solution (solvent not shown). IDRs structures from individual frames at 1-ns intervals were fitted to the R2 segment (helix). Residues along the IDR are shown in blue-to-red colors.

- **Movie SM2: Coarse-grained MD simulations of FAM134B–IDR in solution.**
  The movie spans a 10 $\mu$s coarse-grained MD trajectory of FAM134B-IDR adopting a compact structure in solution (solvent beads not shown; scaled protein-protein interactions using $\alpha = 0.6$). Individual frames at 5-ns intervals were fitted using the R2 segment as a reference. Residues along the IDR are shown in blue-to-red colors.

- **Movie SM3: Coarse-Grained MD simulations of $KALP_{25}$–IDR and RHD–IDR embedded in membranes.**
  The transmembrane $KALP_{25}$-peptide region is shown in blue, and the RHD of FAM134B is shown in green, while the tethered IDR is color-coded in a rainbow scheme. The membrane structure (PO4 beads of POPC lipids) is shown as spheres and colored orange for the upper and blue for the lower leaflet lipids, respectively. The movie (side-views) spans 5 $\mu$s trajectories of $KALP_{25}$–IDR and RHD–IDRs side-by-side, highlighting the comparison of induced local membrane curvature.

- **Movie SM4: IDR collapses and RHD scaffolding of RHD–IDR.**
  The movie shows the trajectory from coarse-grained MD simulations for RHD–IDR (left) in POPC bilayers (not shown), along with its dynamic contact map (right) for the RHD–IDR molecule. The membrane-embedded RHD (green), the tethered IDR (light blue), and the intensity of dynamic RHD–IDR contacts (yellow/orange) are shown along the main chain (side-chain shown for dynamic interactions). The IDR adopts a collapsed and more compact conformation, scaffolding the RHD on the membrane. The movie spans a 1.2 $\mu$s trajectory with snapshots at 2 ns intervals.

- **Movie SM5: Effect of IDRs on curvature sensing.**
  This video presents a side-by-side comparison of curvature sensing simulations of four different protein constructs embedded in buckled bilayers. The membrane-embedded $KALP_{25}$-peptide (blue) and the RHD (green) are embedded in regions of low mean curvature (bottom of the sinusoidal membrane) in the presence and absence of tethered IDRs (rainbow color scheme). Individual lipids are represented by PO4 groups (spheres) and colored independently for upper and lower leaflets (orange/light blue). The movie spans 2 $\mu$s trajectories.

- **Movie S6: IDRs accelerate RHD-mediated protein clustering and spontaneous budding.**
  Representative MD simulations containing nine molecules of $KALP_{25}$-peptides and RHDs with and without tethered IDRs embedded in asymmetric bilayers ($\Delta N = 500$; proteins placed in a square $3 \times 3$ grid 10 nm apart). The membrane-embedded $KALP_{25}$-peptide (blue) and the RHD (green) are embedded in the flat membrane in the presence and absence of tethered IDRs (rainbow color scheme). Individual lipids are represented by PO4 groups (spheres) and colored independently for upper and lower leaflets (orange/light blue). The movies span 5 $\mu$s representative MD trajectories from one of the replicates. IDRs tethered to RHDs mediate IDR-IDR interactions, resulting in the formation of receptor clusters that accelerate the spontaneous budding event.
